## Supplementary Material for "Symbiont loss and gain, rather than co-diversification shapes honeybee gut microbiota diversity and function"

### Supplementary Figures

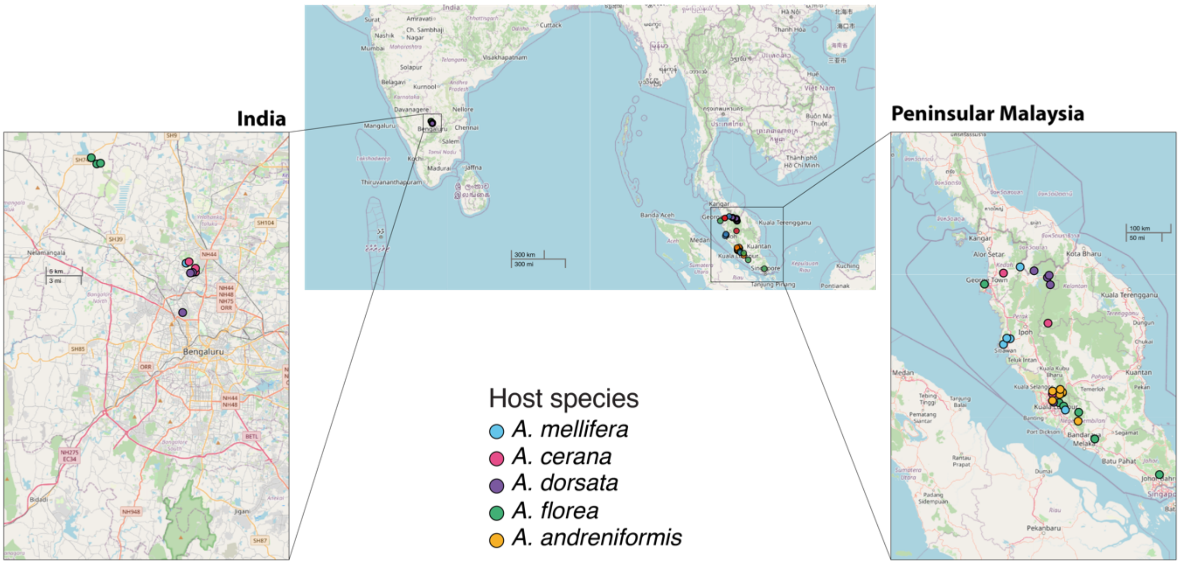


**Supplementary Figure S1. Map of sampling locations.** Colored points indicate the location of the colonies of each host species sampled (size of points are exaggerated for visibility). Map visualization was generated in R Shiny using Leaflet with basemap data from OpenStreetMap (© OpenStreetMap contributors).

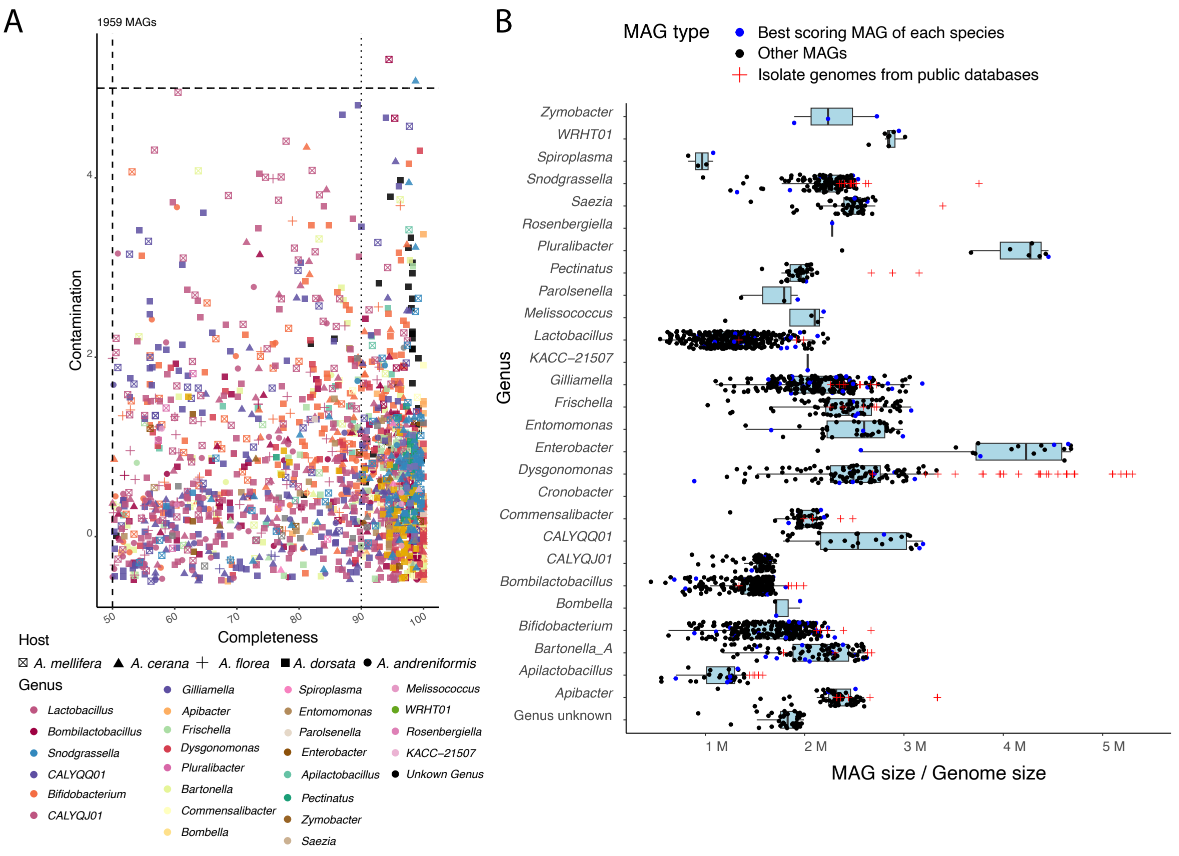


**Supplementary Figure S2. Characteristics of recovered MAGs.** (A) Scatter plot showing the checkM completeness and contamination of each MAG of medium or high quality. Dotted and dashed lines represent the thresholds used to separate high and medium quality MAGs. (B) Boxplot showing the total size of MAGs (sum of length of all contigs binned into the MAG) compared to known sizes of isolated reference genomes of other species from the genus found in NCBI. The best-scoring MAG of each species within the genus is highlighted in blue.

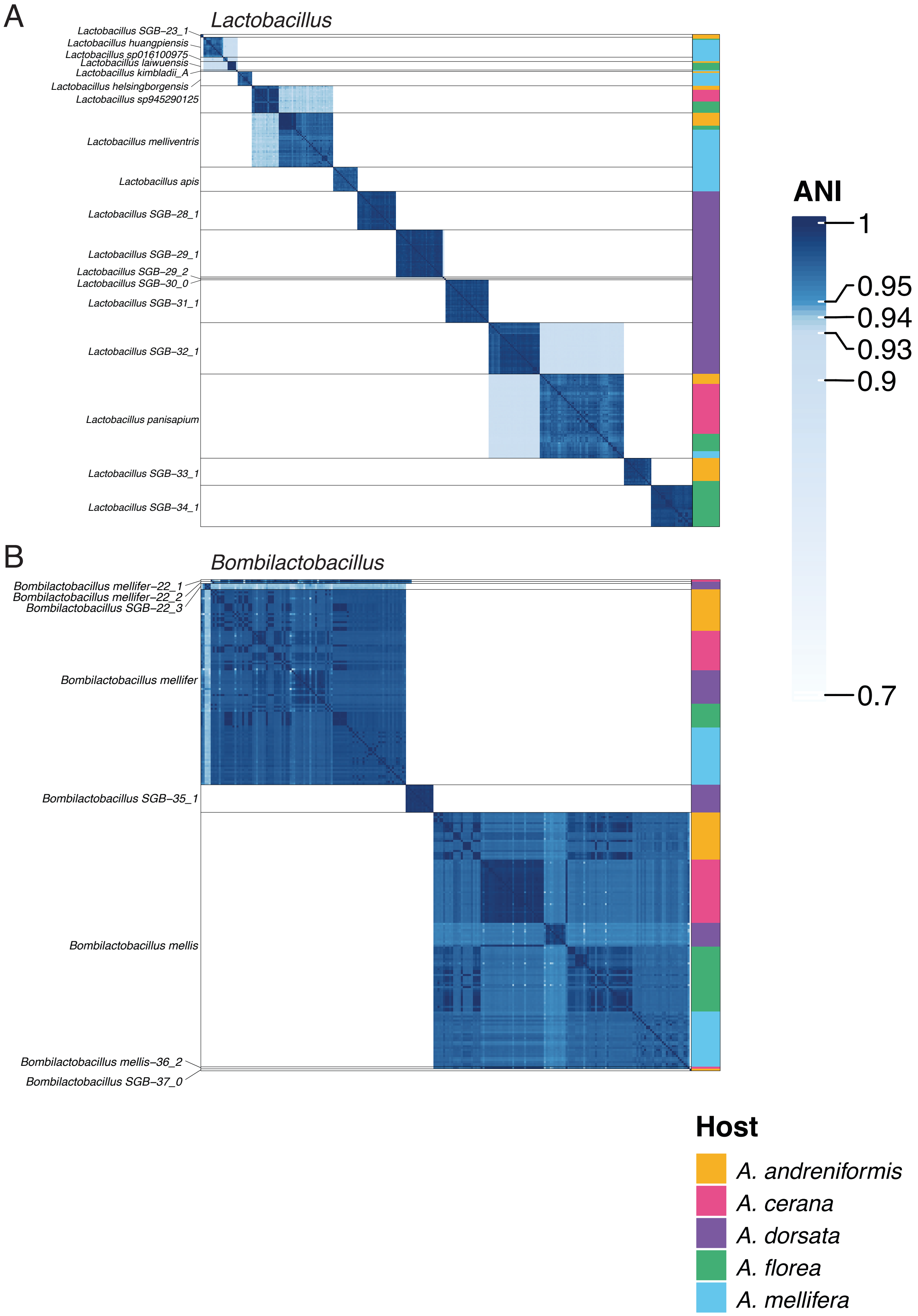

A
Lactobacillus
Lactobacillus SGB–23_1
Lactobacillus huangpiensis
Lactobacillus sp016100975
Lactobacillus laiwuensis
Lactobacillus kimbladii_A
Lactobacillus helsingborgensis
Lactobacillus sp945290125
Lactobacillus melliventris
Lactobacillus apis
Lactobacillus SGB–28_1
Lactobacillus SGB–29_1
Lactobacillus SGB–29_2
Lactobacillus SGB–30_0
Lactobacillus SGB–31_1
Lactobacillus SGB–32_1
Lactobacillus panisapium
Lactobacillus SGB–33_1
Lactobacillus SGB–34_1
B
Bombilactobacillus
Bombilactobacillus mellifer–22_1
Bombilactobacillus mellifer–22_2
Bombilactobacillus SGB–22_3
Bombilactobacillus mellifer
Bombilactobacillus SGB–35_1
Bombilactobacillus mellis
Bombilactobacillus mellis–36_2
Bombilactobacillus SGB–37_0
ANI
1
0.95
0.94
0.93
0.9
0.7
Host
A. andreniformis
A. cerana
A. dorsata
A. florea
A. mellifera

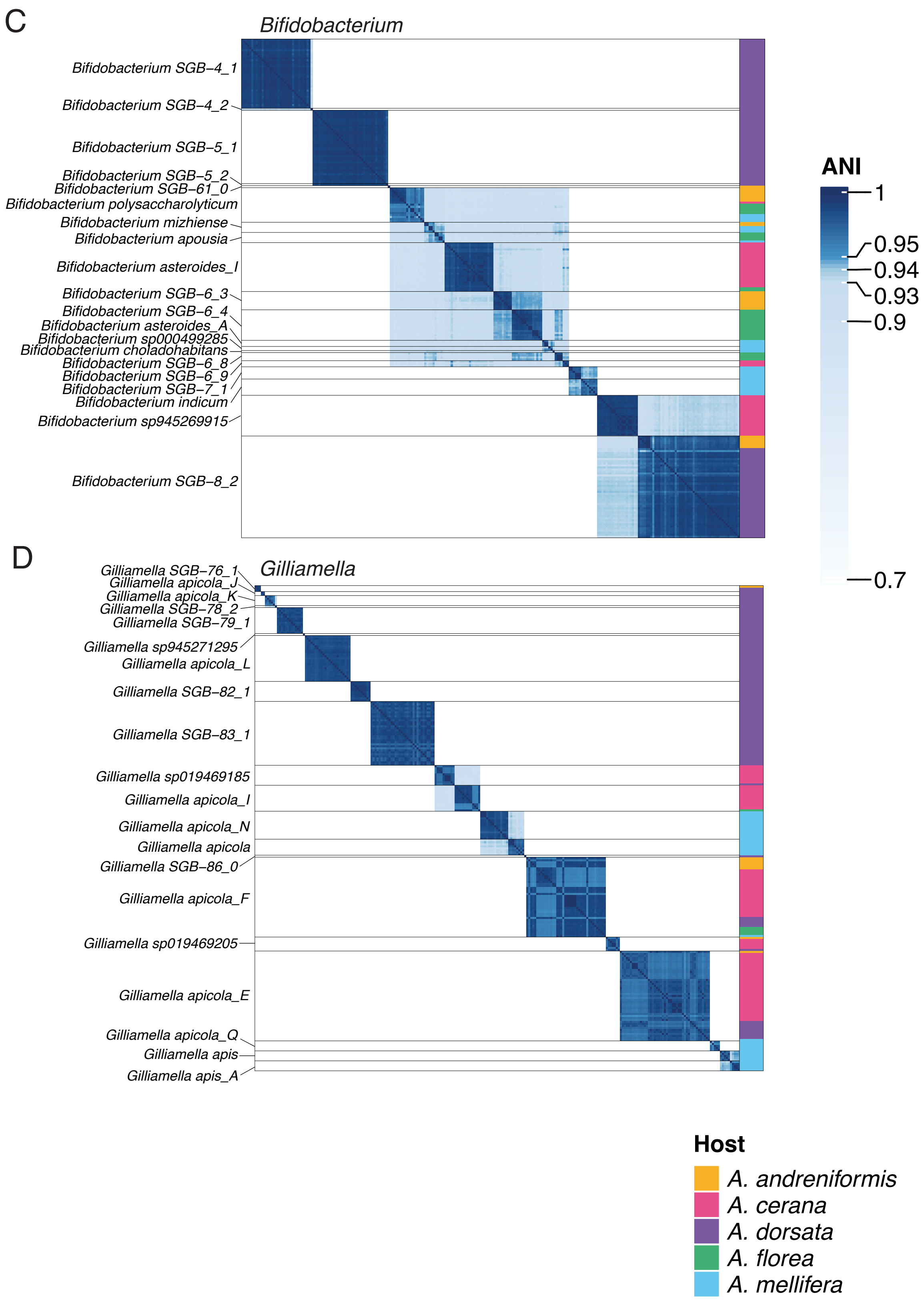
C
Bifidobacterium
Bifidobacterium SGB–4_1
Bifidobacterium SGB–4_2
Bifidobacterium SGB–5_1
Bifidobacterium SGB–5_2
Bifidobacterium SGB–61_0
Bifidobacterium polysaccharolyticum
Bifidobacterium mizhiense
Bifidobacterium apousia
Bifidobacterium asteroides_I
Bifidobacterium SGB–6_3
Bifidobacterium SGB–6_4
Bifidobacterium asteroides_A
Bifidobacterium sp000499285
Bifidobacterium choladohabitans
Bifidobacterium SGB–6_8
Bifidobacterium SGB–6_9
Bifidobacterium SGB–7_1
Bifidobacterium indicum
Bifidobacterium sp945269915
Bifidobacterium SGB–8_2
ANI
1
0.95
0.94
0.93
0.9
0.7
D
Gilliamella
Gilliamella SGB–76_1
Gilliamella apicola_J
Gilliamella apicola_K
Gilliamella SGB–78_2
Gilliamella SGB–79_1
Gilliamella sp945271295
Gilliamella apicola_L
Gilliamella SGB–82_1
Gilliamella SGB–83_1
Gilliamella sp019469185
Gilliamella apicola_I
Gilliamella apicola_N
Gilliamella apicola
Gilliamella SGB–86_0
Gilliamella apicola_F
Gilliamella sp019469205
Gilliamella apicola_E
Gilliamella apicola_Q
Gilliamella apis
Gilliamella apis_A
Host
A. andreniformis
A. cerana
A. dorsata
A. florea
A. mellifera

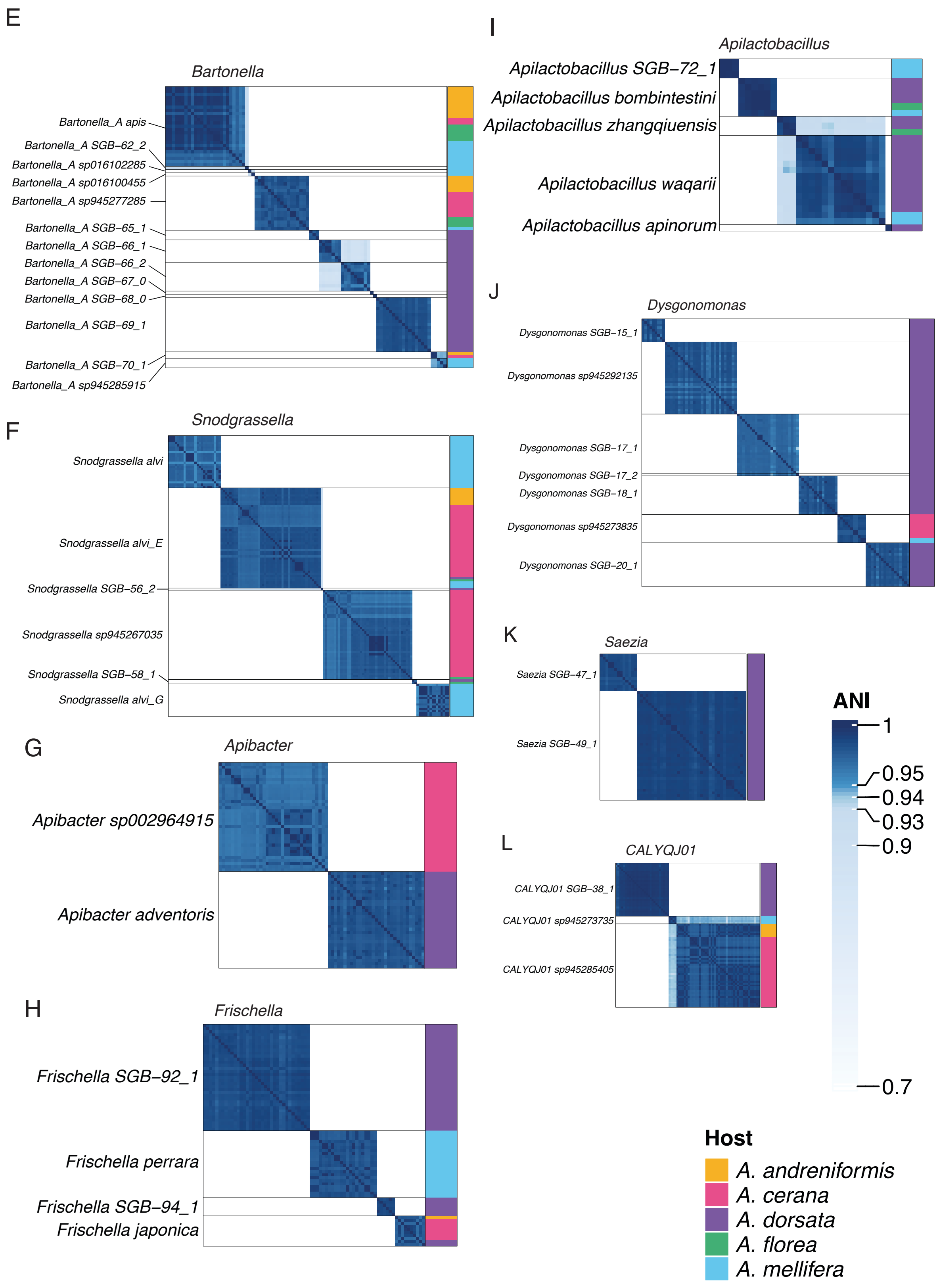

E
Bartonella
Bartonella_A apis
Bartonella_A SGB−62_2
Bartonella_A sp016102285
Bartonella_A sp016100455
Bartonella_A sp945277285
Bartonella_A SGB−65_1
Bartonella_A SGB−66_1
Bartonella_A SGB−66_2
Bartonella_A SGB−67_0
Bartonella_A SGB−68_0
Bartonella_A SGB−69_1
Bartonella_A SGB−70_1
Bartonella_A sp945285915
F
Snodgrassella
Snodgrassella alvi
Snodgrassella alvi_E
Snodgrassella SGB−56_2
Snodgrassella sp945267035
Snodgrassella SGB−58_1
Snodgrassella alvi_G
G
Apibacter
Apibacter sp002964915
Apibacter adventoris
H
Frischella
Frischella SGB−92_1
Frischella perrara
Frischella SGB−94_1
Frischella japonica
I
Apilactobacillus
Apilactobacillus SGB−72_1
Apilactobacillus bombintestini
Apilactobacillus zhangqiuensis
Apilactobacillus waqarii
Apilactobacillus apinorum
J
Dysgonomonas
Dysgonomonas SGB−15_1
Dysgonomonas sp945292135
Dysgonomonas SGB−17_1
Dysgonomonas SGB−17_2
Dysgonomonas SGB−18_1
Dysgonomonas sp945273835
Dysgonomonas SGB−20_1
K
Saezia
Saezia SGB−47_1
Saezia SGB−49_1
L
CALYQJ01
CALYQJ01 SGB−38_1
CALYQJ01 sp945273735
CALYQJ01 sp945285405
ANI
1
0.95
0.94
0.93
0.9
0.7
Host
A. andreniformis
A. cerana
A. dorsata
A. florea
A. mellifera

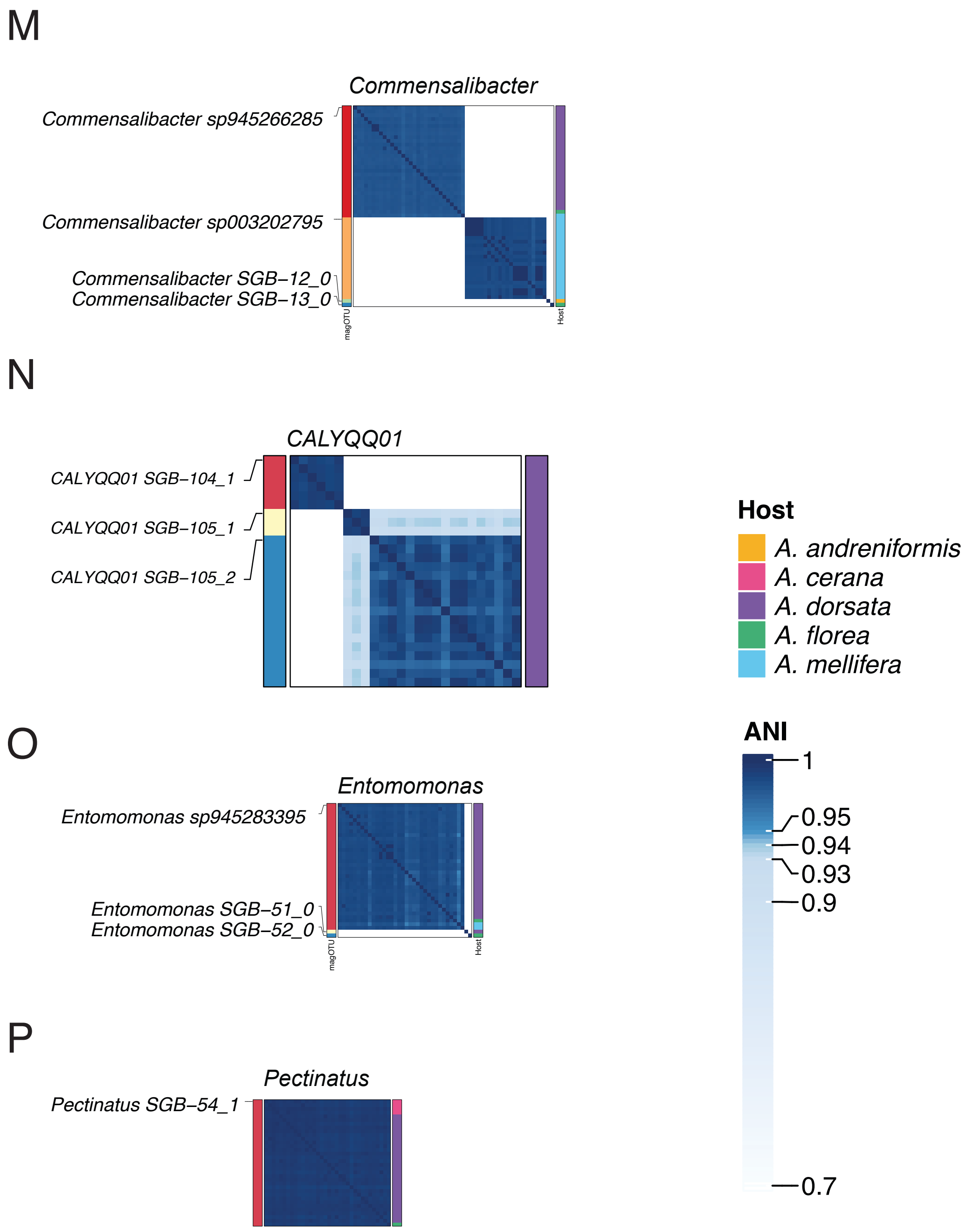
M
Commensalibacter
Commensalibacter sp945266285
Commensalibacter sp003202795
Commensalibacter SGB−12_0
Commensalibacter SGB−13_0
magOTU
Host
N
CALYQQ01
CALYQQ01 SGB−104_1
CALYQQ01 SGB−105_1
CALYQQ01 SGB−105_2
O
Entomomonas
Entomomonas sp945283395
Entomomonas SGB−51_0
Entomomonas SGB−52_0
magOTU
Host
P
Pectinatus
Pectinatus SGB−54_1
Host
A. andreniformis
A. cerana
A. dorsata
A. florea
A. mellifera
ANI
1
0.95
0.94
0.93
0.9
0.7

**Supplementary Figure S3. ANI heatmaps of MAGs.** ANI heatmaps of all medium- and high-quality MAGs per genus with labels on the left indicating the bacterial species assigned and the colors on the right indicating the host species from which the MAG was recovered.

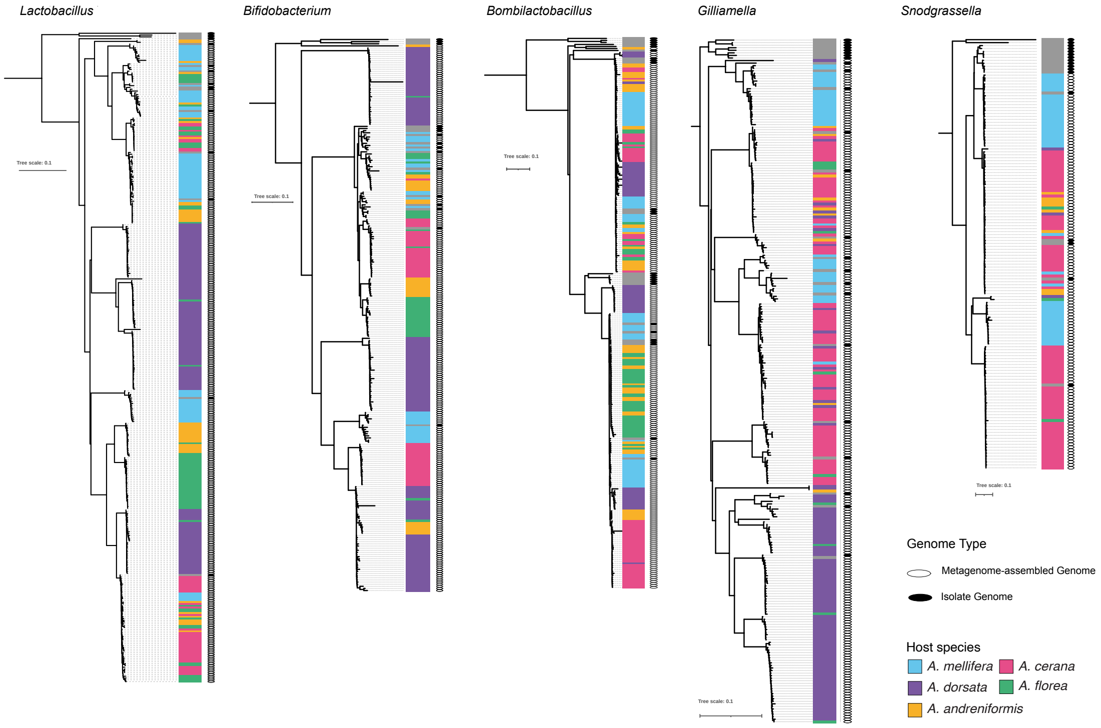


**Supplementary Figure S4. Phylogenies of MAGs and isolate genomes.** Core genome phylogenies, including all high- and medium-quality MAGs of the bacterial genus and selected isolate genomes from the GTDB database for several bacterial species. Tree leaves corresponding to isolate genomes are marked by a black circle. Bacterial species represented by clades of MAGs corresponding to a given harbor isolate genomes of that species within.

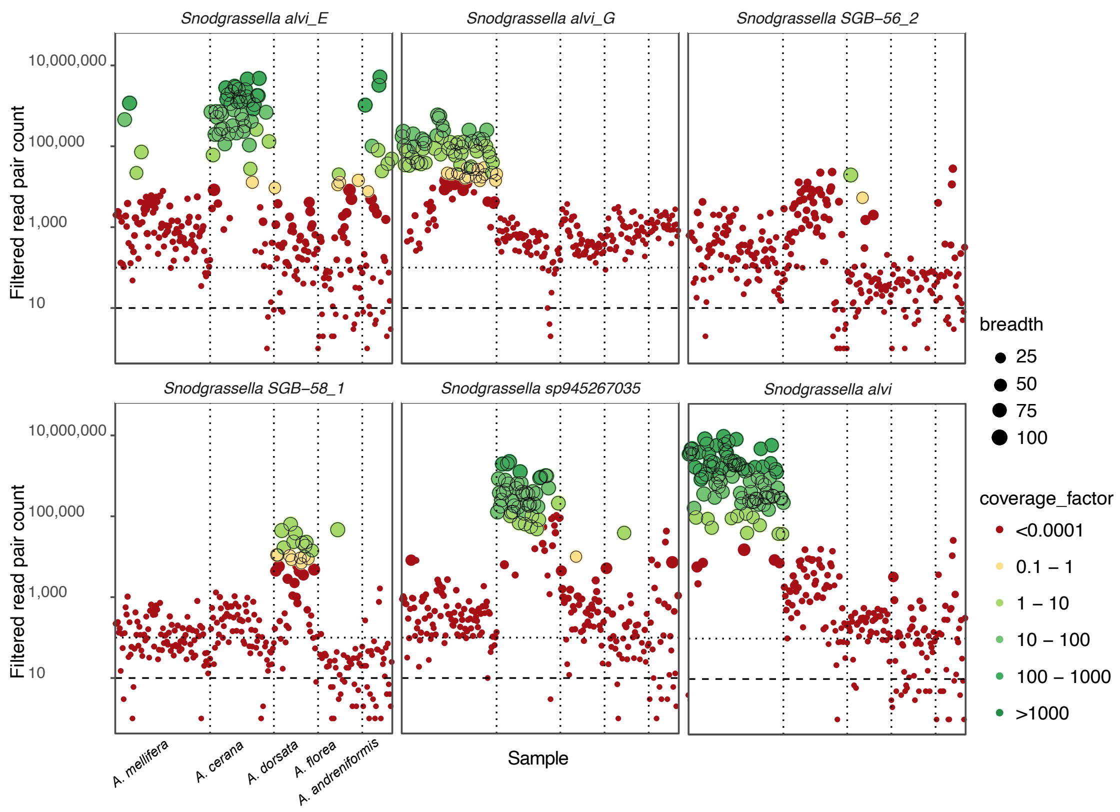


**Supplementary Figure S5. Read mapping to species-representative MAGs.** The panels show an example of six *Snodgrassella* species across samples as an example of read mapping plots for closely related species. Different *Snodgrassella* species are represented in each panel. The x-axis in each panel represents samples grouped by host species (separated by vertical dotted lines), and the y-axis is the number of read-pairs mapping (after filtering to keep only high-quality mapping) to the focal species in each sample. Horizontal dashed and dotted lines represent different depth cut-offs for visualization. The size of the points represents the breadth of the species-representative MAG covered in the sample and the color of the range into which its coverage depth falls. Points outlined with a black line correspond to samples in which the respective species was considered present according to thresholds used in the pipeline.

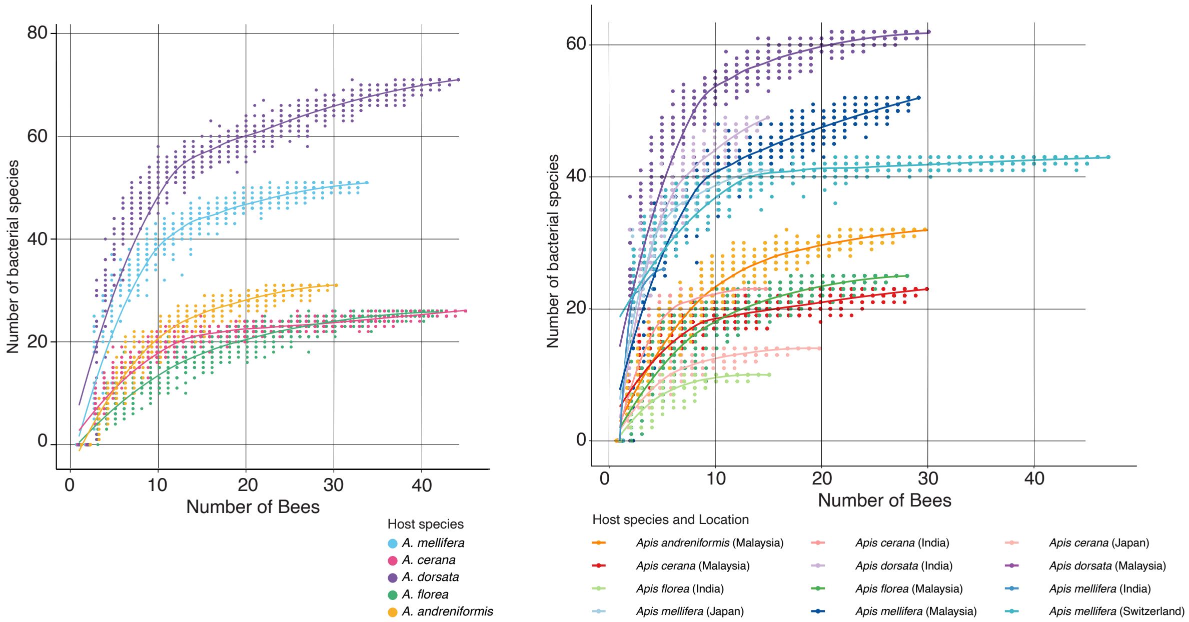


**Supplementary Figure S6. Cumulative curves of the number of bacterial species.**

Number of bacterial species as a function of sample size by host species and sampling location.

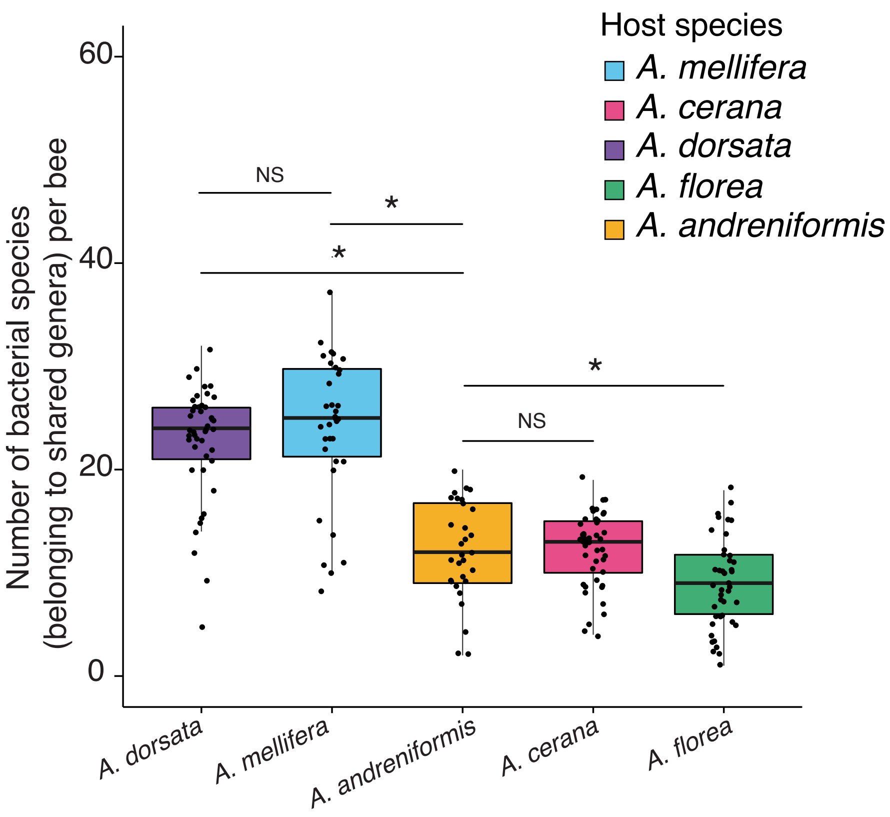


**Supplementary Figure S7. Number of species form shared genera per individual.**

Boxplot showing the number of species only from genera that are detected in at least one bee of all host species (namely, Bombilactobacillus, Lactobacillus, Bifidobacterium, CALYQJ01, Gilliamella, Snodgrassella, Frischella and Bartonella).

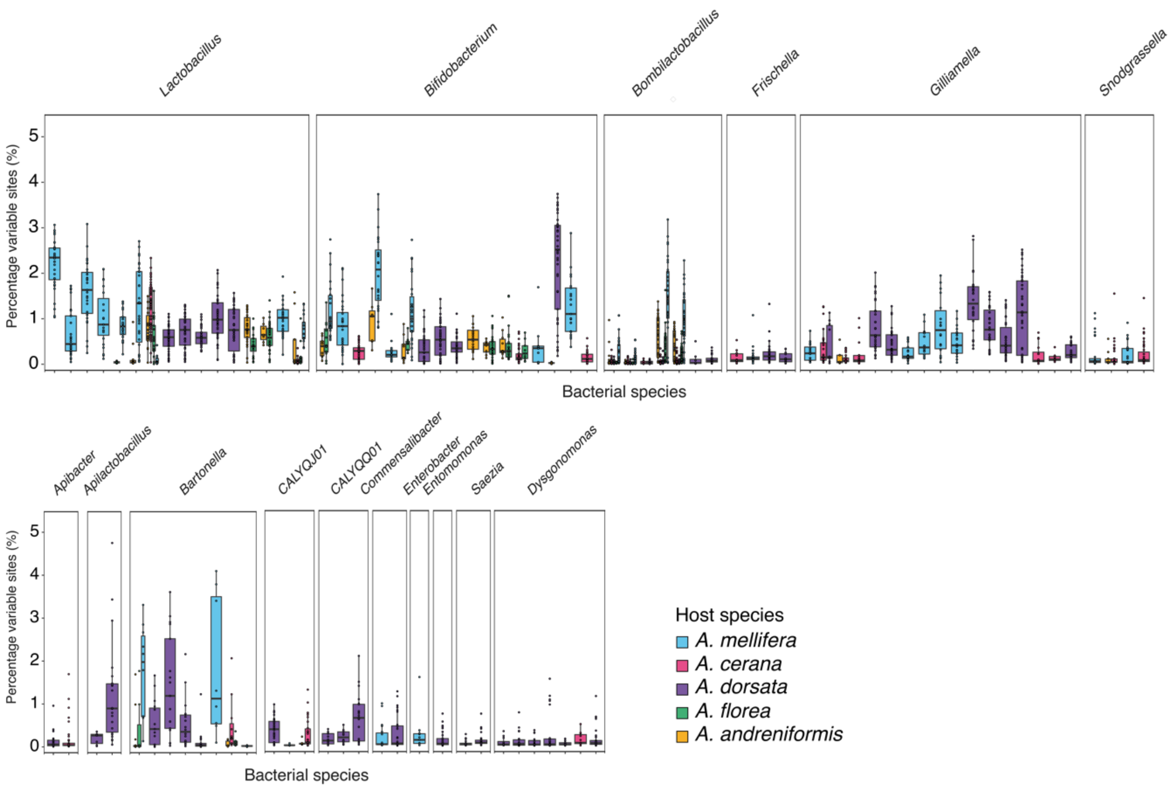


**Supplementary Figure S8. Strain level diversity of each bacterial species across samples.** Boxplots showing the percentage of variable sites (y-axis) marked by SNPs including samples grouped by hosts and for each bacterial species (x-axis).

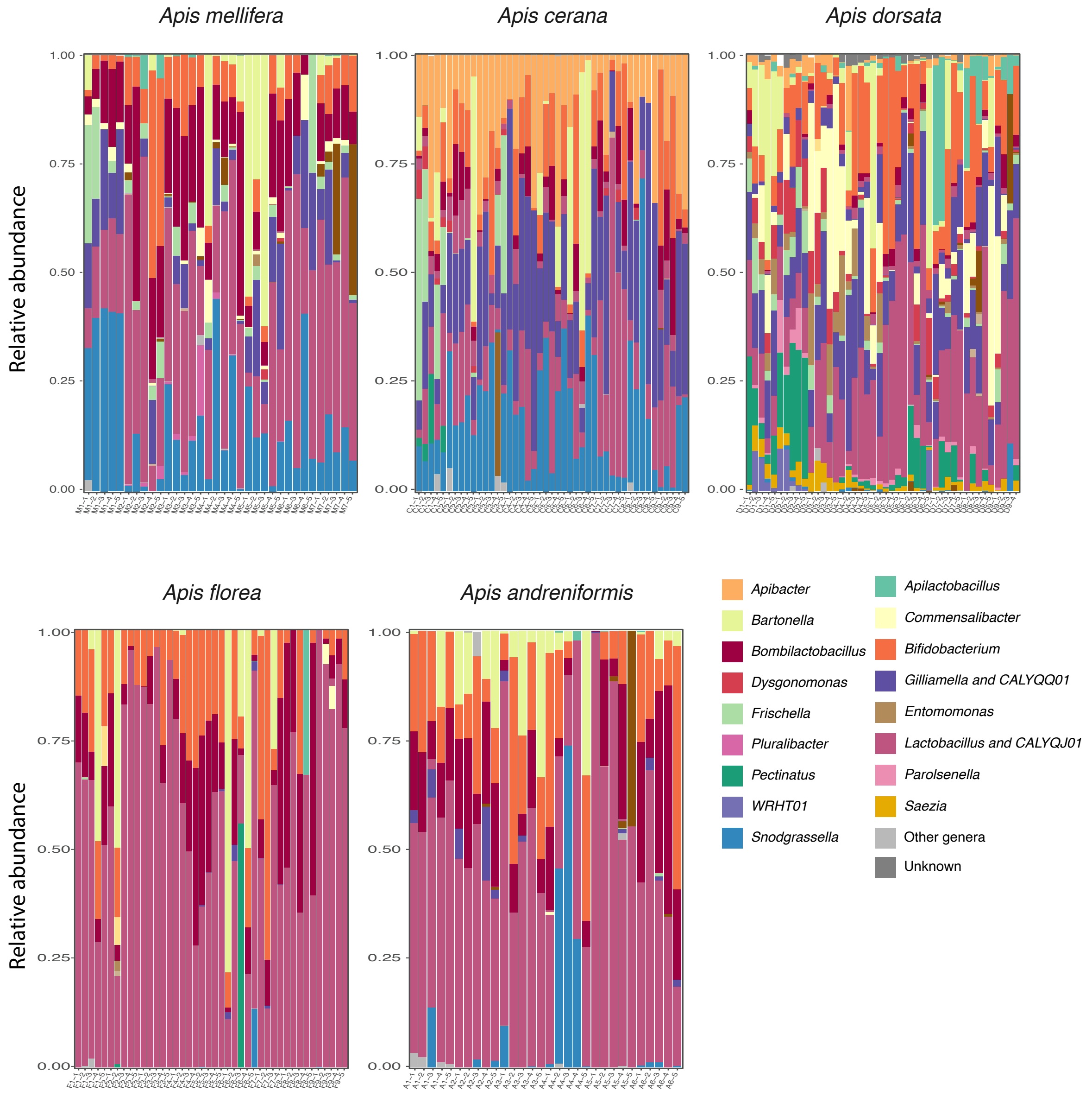


**Supplementary Figure S9. Relative abundance of bacterial genera across samples.** Barplots showing the relative abundance of bacterial genera across samples grouped by host species with white lines within each genus indicating different bacterial species within the genus.

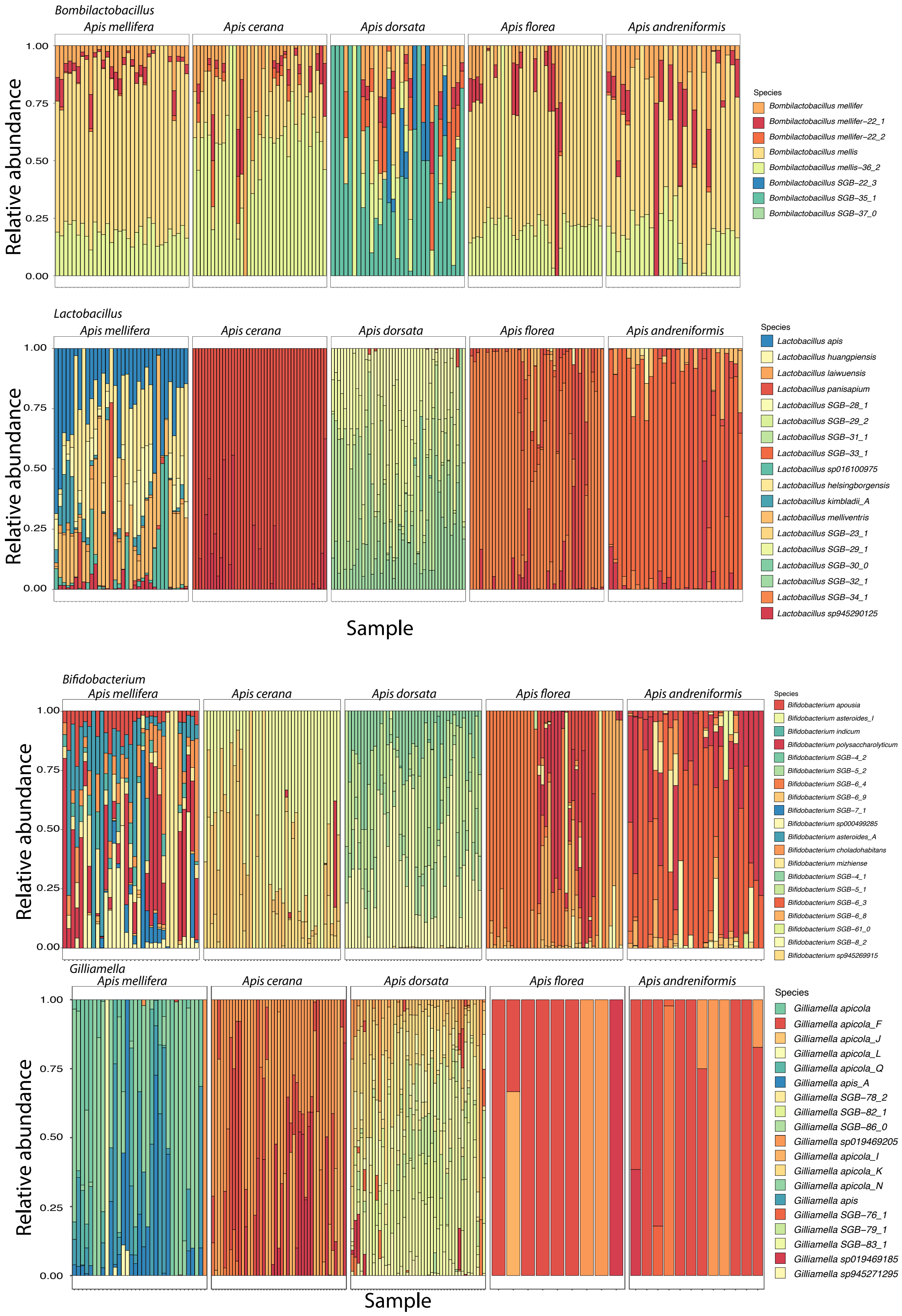
Bombilactobacillus
Apis mellifera
Apis cerana
Apis dorsata
Apis florea
Apis andreniformis
Relative abundance
1.00
0.75
0.50
0.25
0.00
Species
Bombilactobacillus mellifer
Bombilactobacillus mellifer−22_1
Bombilactobacillus mellifer−22_2
Bombilactobacillus mellis
Bombilactobacillus mellis−36_2
Bombilactobacillus SGB−22_3
Bombilactobacillus SGB−35_1
Bombilactobacillus SGB−37_0
Lactobacillus
Species
Lactobacillus apis
Lactobacillus huangpiensis
Lactobacillus laiwuensis
Lactobacillus panisapium
Lactobacillus SGB−28_1
Lactobacillus SGB−29_2
Lactobacillus SGB−31_1
Lactobacillus SGB−33_1
Lactobacillus sp016100975
Lactobacillus helsingborgensis
Lactobacillus kimbladii_A
Lactobacillus melliventris
Lactobacillus SGB−23_1
Lactobacillus SGB−29_1
Lactobacillus SGB−30_0
Lactobacillus SGB−32_1
Lactobacillus SGB−34_1
Lactobacillus sp945290125
Sample
Bifidobacterium
Species
Bifidobacterium apousia
Bifidobacterium asteroides_I
Bifidobacterium indicum
Bifidobacterium polysaccharolyticum
Bifidobacterium SGB−4_2
Bifidobacterium SGB−5_2
Bifidobacterium SGB−6_4
Bifidobacterium SGB−6_9
Bifidobacterium SGB−7_1
Bifidobacterium sp000499285
Bifidobacterium asteroides_A
Bifidobacterium choladohabitans
Bifidobacterium mizhiense
Bifidobacterium SGB−4_1
Bifidobacterium SGB−5_1
Bifidobacterium SGB−6_3
Bifidobacterium SGB−6_8
Bifidobacterium SGB−61_0
Bifidobacterium SGB−8_2
Bifidobacterium sp945269915
Gilliamella
Species
Gilliamella apicola
Gilliamella apicola_F
Gilliamella apicola_J
Gilliamella apicola_L
Gilliamella apicola_Q
Gilliamella apis_A
Gilliamella SGB−78_2
Gilliamella SGB−82_1
Gilliamella SGB−86_0
Gilliamella sp019469205
Gilliamella apicola_I
Gilliamella apicola_K
Gilliamella apicola_N
Gilliamella apis
Gilliamella SGB−76_1
Gilliamella SGB−79_1
Gilliamella SGB−83_1
Gilliamella sp019469185
Gilliamella sp945271295
Sample

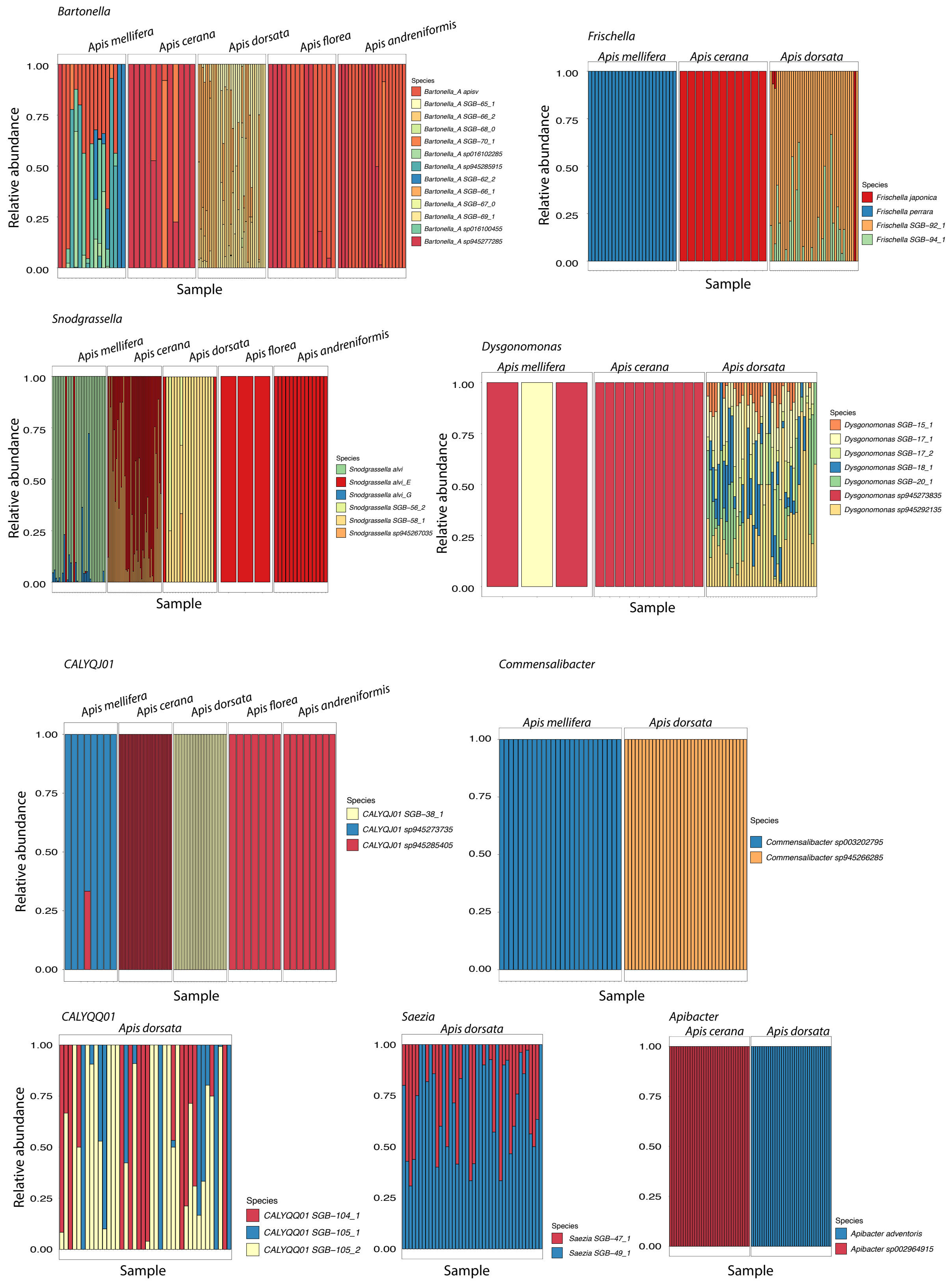


**Supplementary Figure S10. Barplots of the relative abundance of bacterial species by genus.** Barplots summarizing the relative abundance of bacterial species by genus across samples from different host species. Only species found in at least two samples of a given host species and samples that contain at least one bacterial species of the respective genus are included.

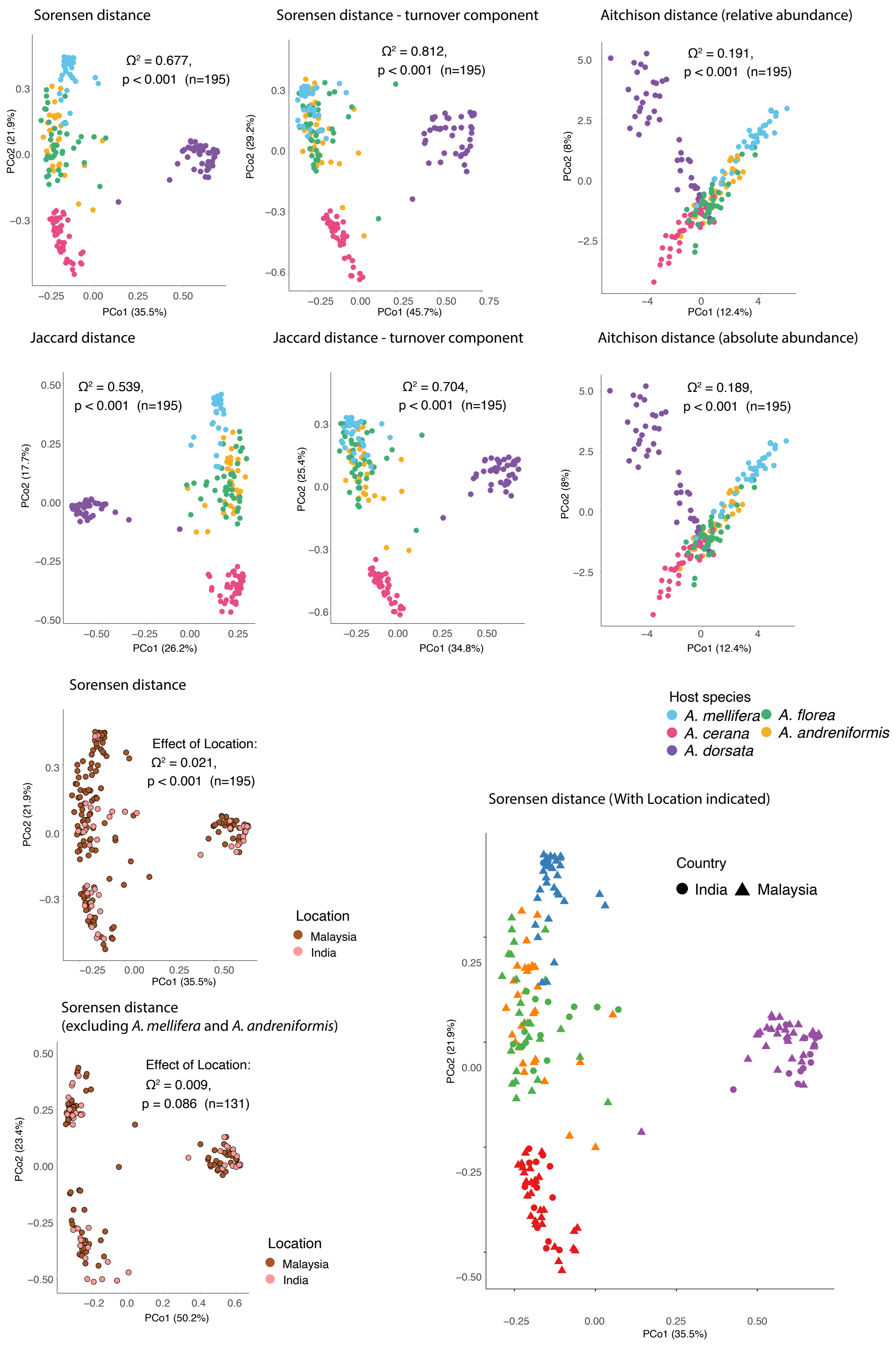


**Supplementary Figure S11. Beta-diversity of bacterial community across host species.**

PCoA of taxonomic diversity using various distance metrics as indicated above each plot.

Results of the PERMANOVA test for the effect of host species are shown on each panel (unless otherwise indicated). For one of the plots by location, two host species that were not sampled equally across locations were excluded to test for the significance of sampling location on the bacterial species diversity more accurately.

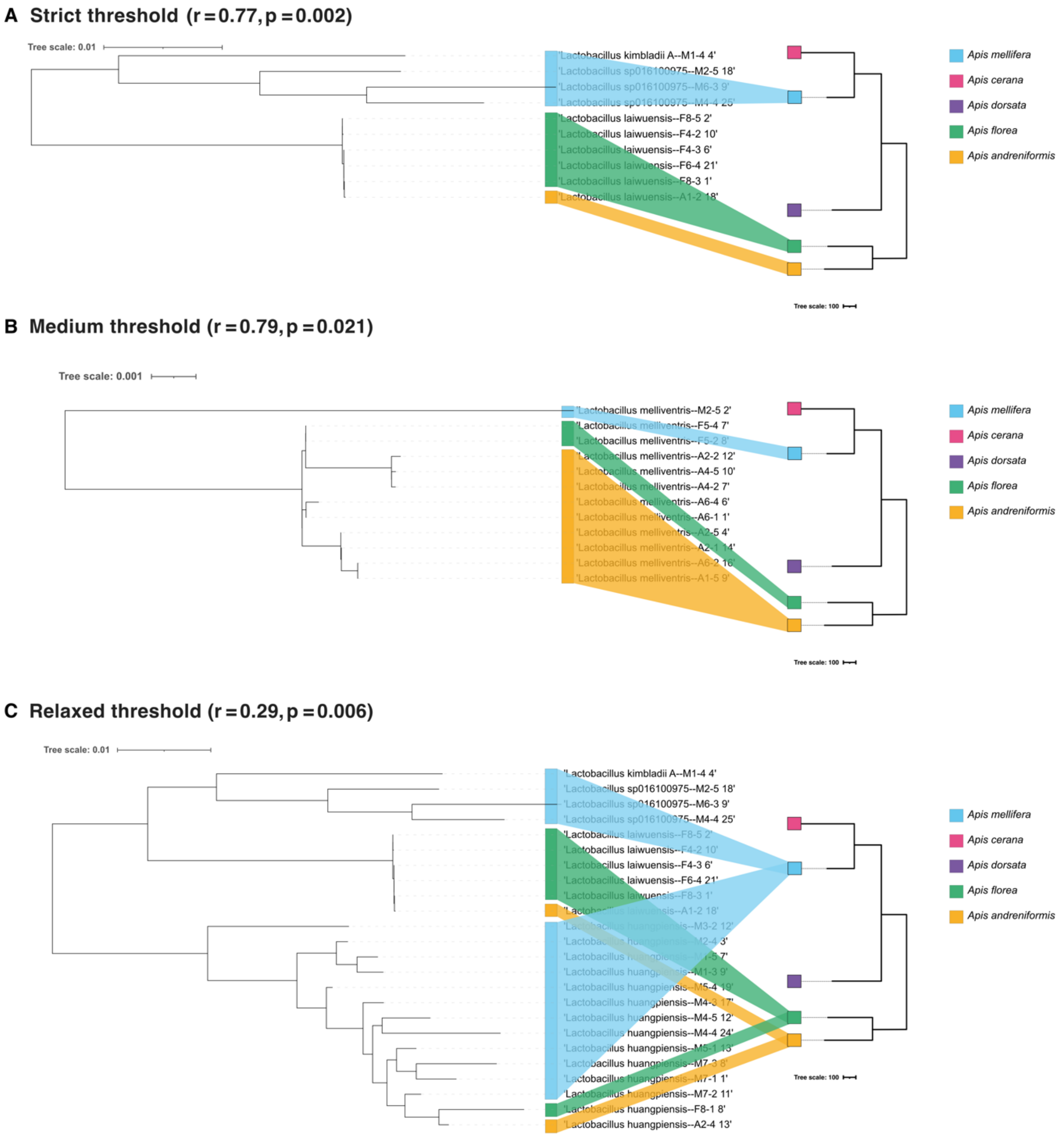

A Strict threshold (r = 0.77, p = 0.002)
Tree scale: 0.01
'Lactobacillus kimbladii A--M1-4 4'
'Lactobacillus sp016100975--M2-5 18'
'Lactobacillus sp016100975--M6-3 9'
'Lactobacillus sp016100975--M4-4 25'
'Lactobacillus laiwuensis--F8-5 2'
'Lactobacillus laiwuensis--F4-2 10'
'Lactobacillus laiwuensis--F4-3 6'
'Lactobacillus laiwuensis--F6-4 21'
'Lactobacillus laiwuensis--F8-3 1'
'Lactobacillus laiwuensis--A1-2 18'
Tree scale: 100
Apis mellifera
Apis cerana
Apis dorsata
Apis florea
Apis andreniformis
B Medium threshold (r = 0.79, p = 0.021)
Tree scale: 0.001
'Lactobacillus melliventris--M2-5 2'
'Lactobacillus melliventris--F5-4 7'
'Lactobacillus melliventris--F5-2 8'
'Lactobacillus melliventris--A2-2 12'
'Lactobacillus melliventris--A4-5 10'
'Lactobacillus melliventris--A4-2 7'
'Lactobacillus melliventris--A6-4 6'
'Lactobacillus melliventris--A6-1 1'
'Lactobacillus melliventris--A2-5 4'
'Lactobacillus melliventris--A2-1 14'
'Lactobacillus melliventris--A6-2 16'
'Lactobacillus melliventris--A1-5 9'
Tree scale: 100
Apis mellifera
Apis cerana
Apis dorsata
Apis florea
Apis andreniformis
C Relaxed threshold (r = 0.29, p = 0.006)
Tree scale: 0.01
'Lactobacillus kimbladii A--M1-4 4'
'Lactobacillus sp016100975--M2-5 18'
'Lactobacillus sp016100975--M6-3 9'
'Lactobacillus sp016100975--M4-4 25'
'Lactobacillus laiwuensis--F8-5 2'
'Lactobacillus laiwuensis--F4-2 10'
'Lactobacillus laiwuensis--F4-3 6'
'Lactobacillus laiwuensis--F6-4 21'
'Lactobacillus laiwuensis--F8-3 1'
'Lactobacillus laiwuensis--A1-2 18'
'Lactobacillus huangpiensis--M3-2 12'
'Lactobacillus huangpiensis--M2-4 3'
'Lactobacillus huangpiensis--M1-5 7'
'Lactobacillus huangpiensis--M1-3 9'
'Lactobacillus huangpiensis--M5-4 19'
'Lactobacillus huangpiensis--M4-3 17'
'Lactobacillus huangpiensis--M4-5 12'
'Lactobacillus huangpiensis--M4-4 24'
'Lactobacillus huangpiensis--M5-1 13'
'Lactobacillus huangpiensis--M7-3 8'
'Lactobacillus huangpiensis--M7-1 1'
'Lactobacillus huangpiensis--M7-2 11'
'Lactobacillus huangpiensis--F8-1 8'
'Lactobacillus huangpiensis--A2-4 13'
Tree scale: 100
Apis mellifera
Apis cerana
Apis dorsata
Apis florea
Apis andreniformis

**D Relaxed threshold ($r = 0.261, p = 0.007$)**

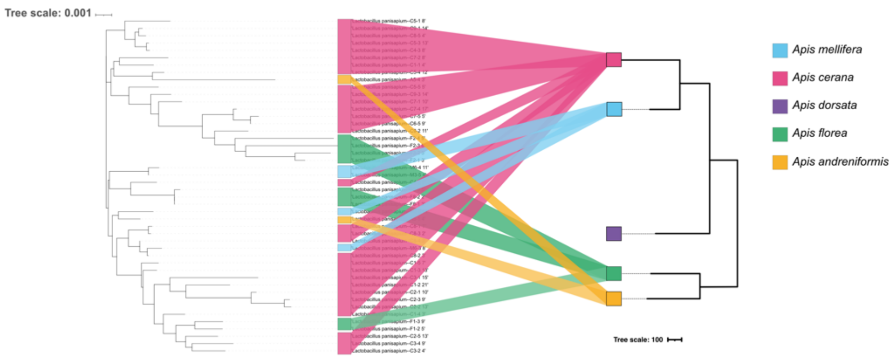


**E Relaxed threshold ($r = 0.496, p = 0.001$)**

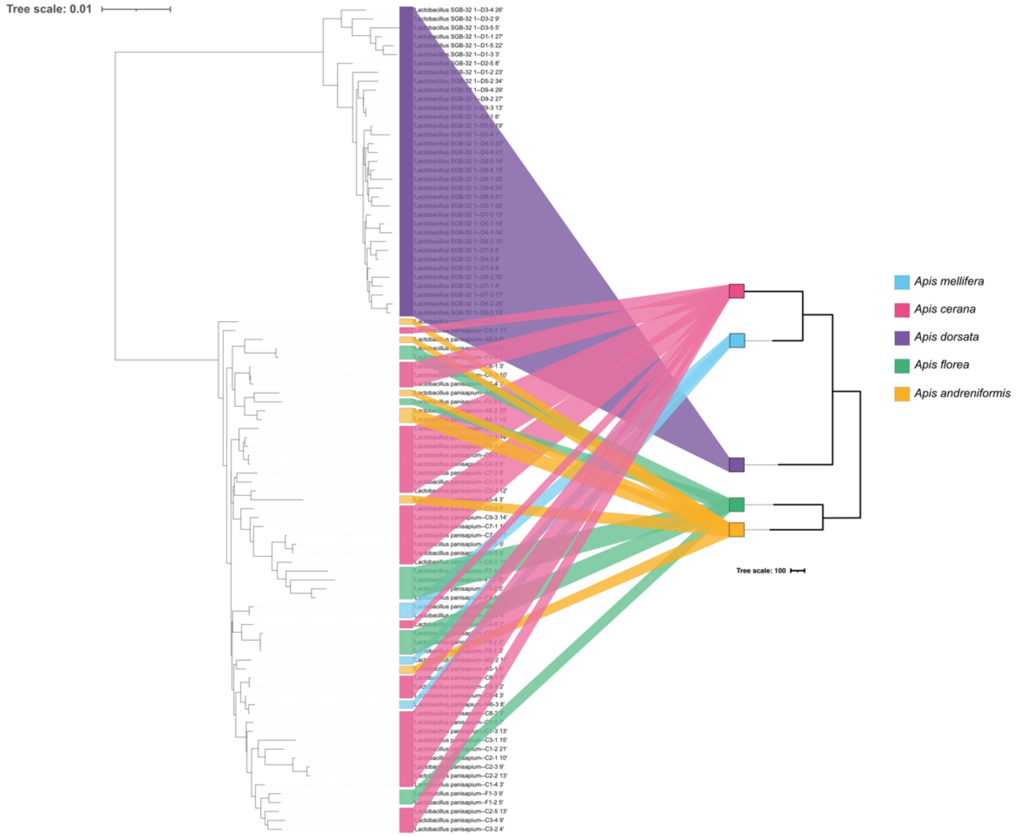


F

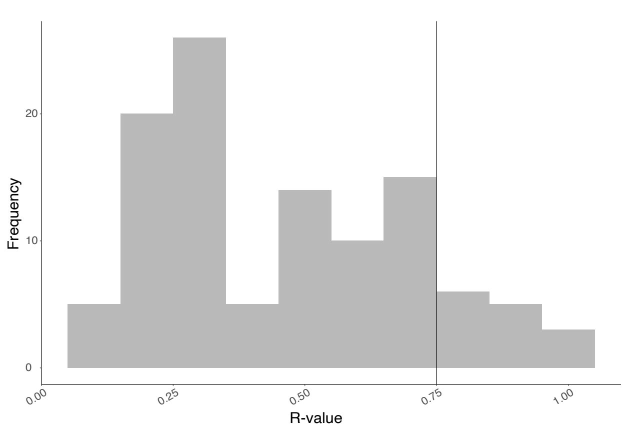

**Supplementary Figure S12. (A-E) Tanglegrams of host (right) and symbiont (left) phylogenies for symbiont nodes showing significant signal for topological congruence at different thresholds (strict, medium, relaxed) and (F) number of significant nodes ($p<0.05$) as a function of R values**. Each tip of the tree is labeled with the corresponding species and MAG name. The colors indicate the host species from which it was recovered. The host tree is based on actual data and to scale according to source (adapted[85]). The host tree scale bar is the inferred number of nucleotide substitutions. The symbiont scale bar represents 0.1 amino-acid substitutions per site.

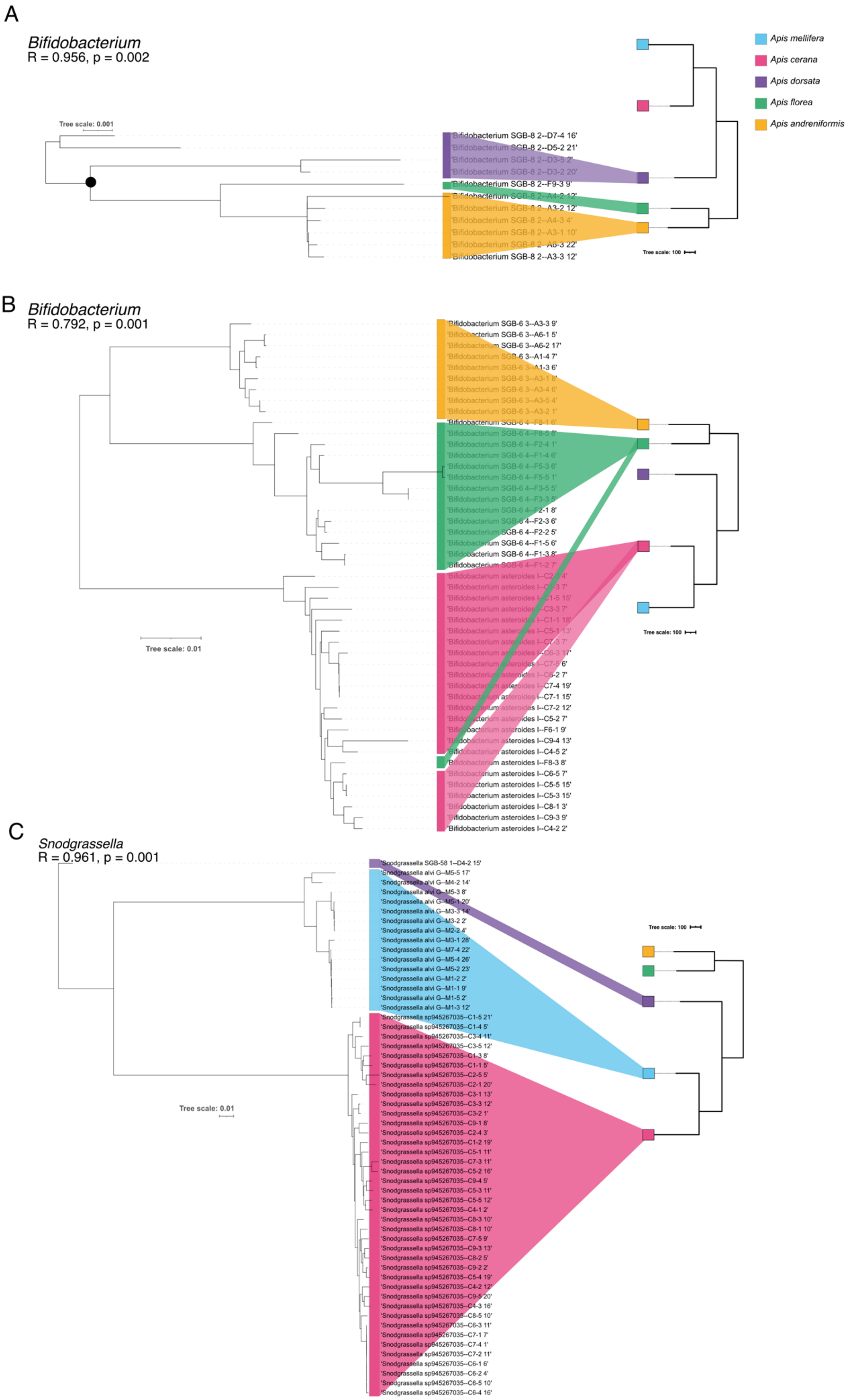

A
Bifidobacterium
R = 0.956, p = 0.002
Tree scale: 0.001
'Bifidobacterium SGB-8 2--D7-4 16'
'Bifidobacterium SGB-8 2--D5-2 21'
'Bifidobacterium SGB-8 2--D3-5 2'
'Bifidobacterium SGB-8 2--D3-2 20'
'Bifidobacterium SGB-8 2--F9-3 9'
'Bifidobacterium SGB-8 2--A4-2 12'
'Bifidobacterium SGB-8 2--A3-2 12'
'Bifidobacterium SGB-8 2--A4-3 4'
'Bifidobacterium SGB-8 2--A3-1 10'
'Bifidobacterium SGB-8 2--A6-3 22'
'Bifidobacterium SGB-8 2--A3-3 12'
Tree scale: 100
Apis mellifera
Apis cerana
Apis dorsata
Apis florea
Apis andreniformis
B
Bifidobacterium
R = 0.792, p = 0.001
'Bifidobacterium SGB-6 3--A3-3 9'
'Bifidobacterium SGB-6 3--A6-1 5'
'Bifidobacterium SGB-6 3--A6-2 17'
'Bifidobacterium SGB-6 3--A1-4 7'
'Bifidobacterium SGB-6 3--A1-3 6'
'Bifidobacterium SGB-6 3--A3-1 8'
'Bifidobacterium SGB-6 3--A3-4 6'
'Bifidobacterium SGB-6 3--A3-5 4'
'Bifidobacterium SGB-6 3--A3-2 1'
'Bifidobacterium SGB-6 4--F8-1 6'
'Bifidobacterium SGB-6 4--F8-5 8'
'Bifidobacterium SGB-6 4--F2-4 1'
'Bifidobacterium SGB-6 4--F1-4 6'
'Bifidobacterium SGB-6 4--F5-3 6'
'Bifidobacterium SGB-6 4--F5-5 1'
'Bifidobacterium SGB-6 4--F3-5 5'
'Bifidobacterium SGB-6 4--F3-3 5'
'Bifidobacterium SGB-6 4--F2-1 8'
'Bifidobacterium SGB-6 4--F2-3 6'
'Bifidobacterium SGB-6 4--F2-2 5'
'Bifidobacterium SGB-6 4--F1-5 6'
'Bifidobacterium SGB-6 4--F1-3 8'
'Bifidobacterium SGB-6 4--F1-2 7'
'Bifidobacterium asteroides I--C2-4 4'
'Bifidobacterium asteroides I--C6-3 7'
'Bifidobacterium asteroides I--C1-5 15'
'Bifidobacterium asteroides I--C3-3 7'
'Bifidobacterium asteroides I--C1-1 18'
'Bifidobacterium asteroides I--C5-1 13'
'Bifidobacterium asteroides I--C7-3 7'
'Bifidobacterium asteroides I--C6-3 17'
'Bifidobacterium asteroides I--C7-5 6'
'Bifidobacterium asteroides I--C6-2 7'
'Bifidobacterium asteroides I--C7-4 19'
'Bifidobacterium asteroides I--C7-1 15'
'Bifidobacterium asteroides I--C7-2 12'
'Bifidobacterium asteroides I--C5-2 7'
'Bifidobacterium asteroides I--F6-1 9'
'Bifidobacterium asteroides I--C9-4 13'
'Bifidobacterium asteroides I--C4-5 2'
'Bifidobacterium asteroides I--F8-3 8'
'Bifidobacterium asteroides I--C6-5 7'
'Bifidobacterium asteroides I--C5-5 15'
'Bifidobacterium asteroides I--C5-3 15'
'Bifidobacterium asteroides I--C8-1 3'
'Bifidobacterium asteroides I--C9-3 9'
'Bifidobacterium asteroides I--C4-2 2'
Tree scale: 0.01
Tree scale: 100
C
Snodgrassella
R = 0.961, p = 0.001
'Snodgrassella SGB-58 1--D4-2 15'
'Snodgrassella alvi G--M5-5 17'
'Snodgrassella alvi G--M4-2 14'
'Snodgrassella alvi G--M5-3 8'
'Snodgrassella alvi G--M5-1 20'
'Snodgrassella alvi G--M3-3 14'
'Snodgrassella alvi G--M3-2 2'
'Snodgrassella alvi G--M2-2 4'
'Snodgrassella alvi G--M3-1 28'
'Snodgrassella alvi G--M7-4 22'
'Snodgrassella alvi G--M5-4 26'
'Snodgrassella alvi G--M5-2 23'
'Snodgrassella alvi G--M1-2 2'
'Snodgrassella alvi G--M1-1 9'
'Snodgrassella alvi G--M1-5 2'
'Snodgrassella alvi G--M1-3 12'
'Snodgrassella sp945267035--C1-5 21'
'Snodgrassella sp945267035--C1-4 5'
'Snodgrassella sp945267035--C3-4 11'
'Snodgrassella sp945267035--C3-5 12'
'Snodgrassella sp945267035--C1-3 8'
'Snodgrassella sp945267035--C1-1 5'
'Snodgrassella sp945267035--C2-5 5'
'Snodgrassella sp945267035--C2-1 20'
'Snodgrassella sp945267035--C3-1 13'
'Snodgrassella sp945267035--C3-3 12'
'Snodgrassella sp945267035--C3-2 1'
'Snodgrassella sp945267035--C9-1 8'
'Snodgrassella sp945267035--C2-4 3'
'Snodgrassella sp945267035--C1-2 19'
'Snodgrassella sp945267035--C5-1 11'
'Snodgrassella sp945267035--C7-3 11'
'Snodgrassella sp945267035--C5-2 16'
'Snodgrassella sp945267035--C9-4 5'
'Snodgrassella sp945267035--C5-3 11'
'Snodgrassella sp945267035--C5-5 12'
'Snodgrassella sp945267035--C4-1 2'
'Snodgrassella sp945267035--C8-3 10'
'Snodgrassella sp945267035--C8-1 10'
'Snodgrassella sp945267035--C7-5 9'
'Snodgrassella sp945267035--C9-3 13'
'Snodgrassella sp945267035--C8-2 5'
'Snodgrassella sp945267035--C9-2 2'
'Snodgrassella sp945267035--C5-4 19'
'Snodgrassella sp945267035--C4-2 12'
'Snodgrassella sp945267035--C9-5 20'
'Snodgrassella sp945267035--C4-3 16'
'Snodgrassella sp945267035--C8-5 10'
'Snodgrassella sp945267035--C6-3 11'
'Snodgrassella sp945267035--C7-1 7'
'Snodgrassella sp945267035--C7-4 1'
'Snodgrassella sp945267035--C7-2 11'
'Snodgrassella sp945267035--C6-1 6'
'Snodgrassella sp945267035--C6-2 4'
'Snodgrassella sp945267035--C6-5 10'
'Snodgrassella sp945267035--C6-4 16'
Tree scale: 100
Tree scale: 0.01

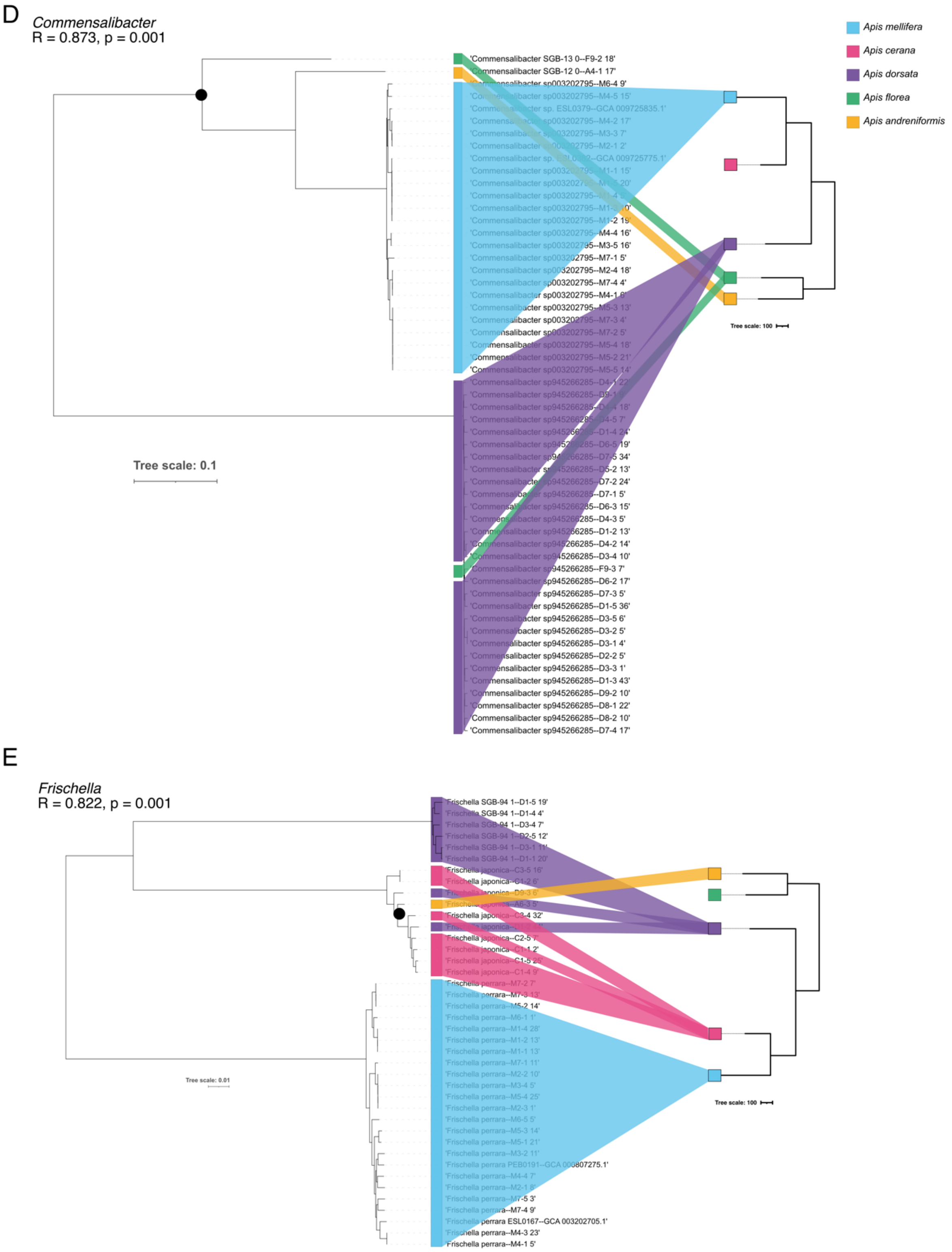
D
Commensalibacter
R = 0.873, p = 0.001
'Commensalibacter SGB-13 0--F9-2 18'
'Commensalibacter SGB-12 0--A4-1 17'
'Commensalibacter sp003202795--M6-4 9'
'Commensalibacter sp003202795--M4-5 15'
'Commensalibacter sp. ESL0379--GCA 009725835.1'
'Commensalibacter sp003202795--M4-2 17'
'Commensalibacter sp003202795--M3-3 7'
'Commensalibacter sp003202795--M2-1 2'
'Commensalibacter sp. ESL0382--GCA 009725775.1'
'Commensalibacter sp003202795--M1-1 15'
'Commensalibacter sp003202795--M1-5 20'
'Commensalibacter sp003202795--M1-4 5'
'Commensalibacter sp003202795--M1-3 10'
'Commensalibacter sp003202795--M1-2 19'
'Commensalibacter sp003202795--M4-4 16'
'Commensalibacter sp003202795--M3-5 16'
'Commensalibacter sp003202795--M7-1 5'
'Commensalibacter sp003202795--M2-4 18'
'Commensalibacter sp003202795--M7-4 4'
'Commensalibacter sp003202795--M4-1 6'
'Commensalibacter sp003202795--M5-3 13'
'Commensalibacter sp003202795--M7-3 4'
'Commensalibacter sp003202795--M7-2 5'
'Commensalibacter sp003202795--M5-4 18'
'Commensalibacter sp003202795--M5-2 21'
'Commensalibacter sp003202795--M5-5 14'
'Commensalibacter sp945266285--D4-1 22'
'Commensalibacter sp945266285--D9-1 9'
'Commensalibacter sp945266285--D4-4 18'
'Commensalibacter sp945266285--D4-5 7'
'Commensalibacter sp945266285--D1-4 24'
'Commensalibacter sp945266285--D6-5 19'
'Commensalibacter sp945266285--D7-5 34'
'Commensalibacter sp945266285--D5-2 13'
'Commensalibacter sp945266285--D7-2 24'
'Commensalibacter sp945266285--D7-1 5'
'Commensalibacter sp945266285--D6-3 15'
'Commensalibacter sp945266285--D4-3 5'
'Commensalibacter sp945266285--D1-2 13'
'Commensalibacter sp945266285--D4-2 14'
'Commensalibacter sp945266285--D3-4 10'
'Commensalibacter sp945266285--F9-3 7'
'Commensalibacter sp945266285--D6-2 17'
'Commensalibacter sp945266285--D7-3 5'
'Commensalibacter sp945266285--D1-5 36'
'Commensalibacter sp945266285--D3-5 6'
'Commensalibacter sp945266285--D3-2 5'
'Commensalibacter sp945266285--D3-1 4'
'Commensalibacter sp945266285--D2-2 5'
'Commensalibacter sp945266285--D3-3 1'
'Commensalibacter sp945266285--D1-3 43'
'Commensalibacter sp945266285--D9-2 10'
'Commensalibacter sp945266285--D8-1 22'
'Commensalibacter sp945266285--D8-2 10'
'Commensalibacter sp945266285--D7-4 17'
Tree scale: 0.1
Tree scale: 100
Apis mellifera
Apis cerana
Apis dorsata
Apis florea
Apis andreniformis
E
Frischella
R = 0.822, p = 0.001
'Frischella SGB-94 1--D1-5 19'
'Frischella SGB-94 1--D1-4 4'
'Frischella SGB-94 1--D3-4 7'
'Frischella SGB-94 1--D2-5 12'
'Frischella SGB-94 1--D3-1 11'
'Frischella SGB-94 1--D1-1 20'
'Frischella japonica--C3-5 16'
'Frischella japonica--C1-2 6'
'Frischella japonica--D9-3 6'
'Frischella japonica--A6-3 5'
'Frischella japonica--C3-4 32'
'Frischella japonica--D1-2 44'
'Frischella japonica--C2-5 7'
'Frischella japonica--C1-1 2'
'Frischella japonica--C1-5 25'
'Frischella japonica--C1-4 9'
'Frischella perrara--M7-2 7'
'Frischella perrara--M7-3 13'
'Frischella perrara--M5-2 14'
'Frischella perrara--M6-1 1'
'Frischella perrara--M1-4 28'
'Frischella perrara--M1-2 13'
'Frischella perrara--M1-1 13'
'Frischella perrara--M7-1 11'
'Frischella perrara--M2-2 10'
'Frischella perrara--M3-4 5'
'Frischella perrara--M5-4 25'
'Frischella perrara--M2-3 1'
'Frischella perrara--M6-5 5'
'Frischella perrara--M5-3 14'
'Frischella perrara--M5-1 21'
'Frischella perrara--M3-2 11'
'Frischella perrara PEB0191--GCA 000807275.1'
'Frischella perrara--M4-4 7'
'Frischella perrara--M2-1 8'
'Frischella perrara--M7-5 3'
'Frischella perrara--M7-4 9'
'Frischella perrara ESL0167--GCA 003202705.1'
'Frischella perrara--M4-3 23'
'Frischella perrara--M4-1 5'
Tree scale: 0.01
Tree scale: 100

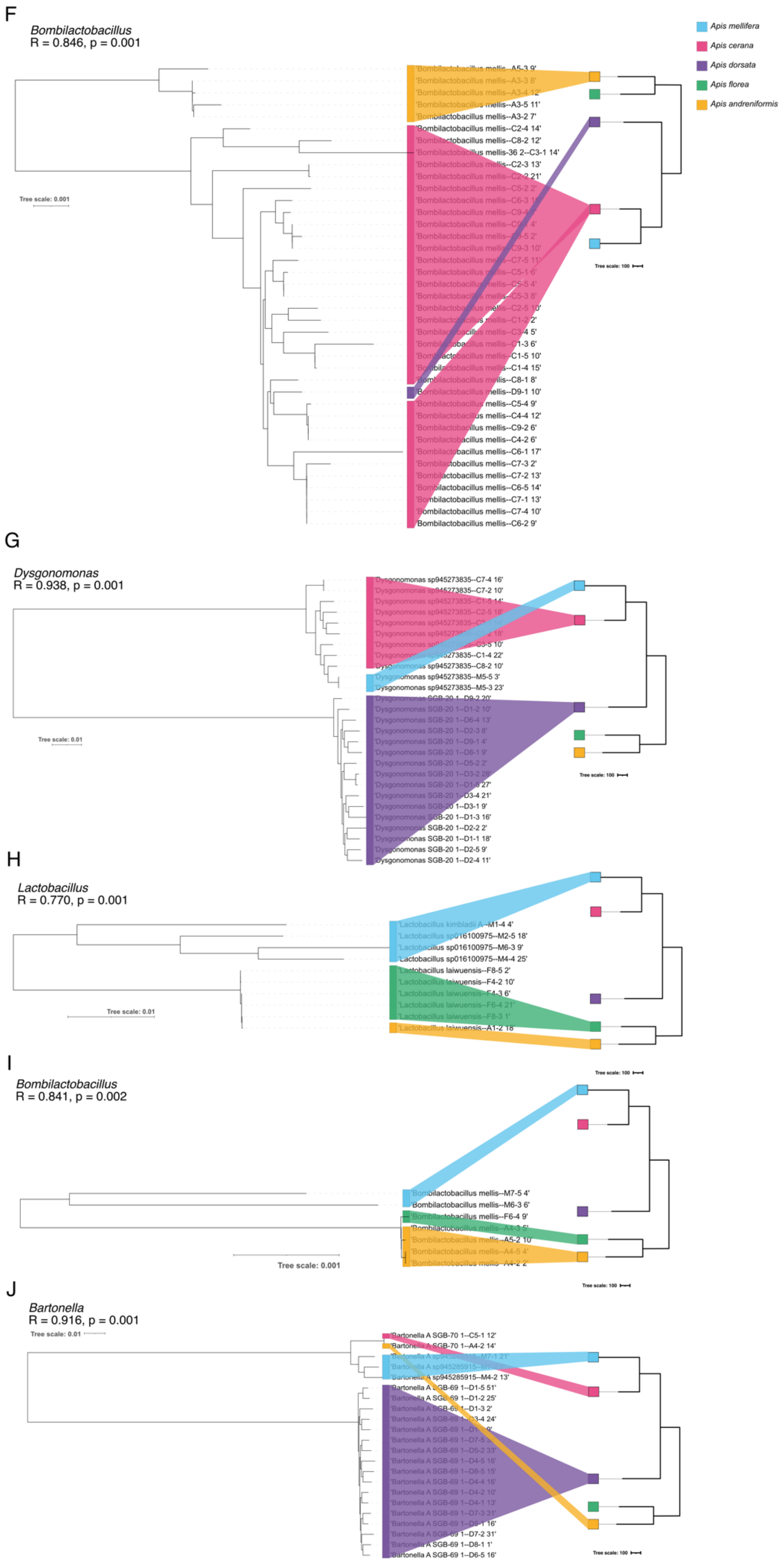

F
Bombilactobacillus
R = 0.846, p = 0.001
Apis mellifera
Apis cerana
Apis dorsata
Apis florea
Apis andreniformis
Tree scale: 0.001
Tree scale: 100
G
Dysgonomonas
R = 0.938, p = 0.001
Tree scale: 0.01
Tree scale: 100
H
Lactobacillus
R = 0.770, p = 0.001
Tree scale: 0.01
I
Bombilactobacillus
R = 0.841, p = 0.002
Tree scale: 100
Tree scale: 0.001
Tree scale: 100
J
Bartonella
R = 0.916, p = 0.001
Tree scale: 0.01
Tree scale: 100

**Supplementary Figure S13. Tanglegrams of host (right) and symbiont (left) phylogenies for symbiont nodes showing significant signal for topological congruence ($R > 0.75$, $p<0.01$)**. These trees correspond to the subtrees of the nodes indicated with a black filled circle in **Fig. 3** in the main text. In three cases, a significant node was nested within the depicted subtrees (see black circle in panel A, D and E). Each tip of the tree is labeled with the corresponding species and MAG name. The colors indicate the host species from which it was recovered. The host tree is based on actual data and to scale according to source (adapted[85]). The host tree scale bar is the inferred number of nucleotide substitutions. The symbiont scale bar represents 0.1 amino-acid substitutions per site. In contrast to the trees shown in panel A-D, which show clear evidence of co-diversification, subtrees in panel E and F are not congruent with the host tree topology, and in the subtrees in panel G-J , some branches between symbionts from different hosts are not consistent with host divergence. For example, there is little to no divergence between MAGs from *A. florea* and *A. andreniformis* in the subtrees of panel H and I, despite the fact that these two bee species have diverged 6 mya. Also there is little divergence between MAGs from *A. mellifera* and *A. cerana* in subtrees in panel G and J relatively to the very deeply branching sister clade of *A. dorsata* MAGs.

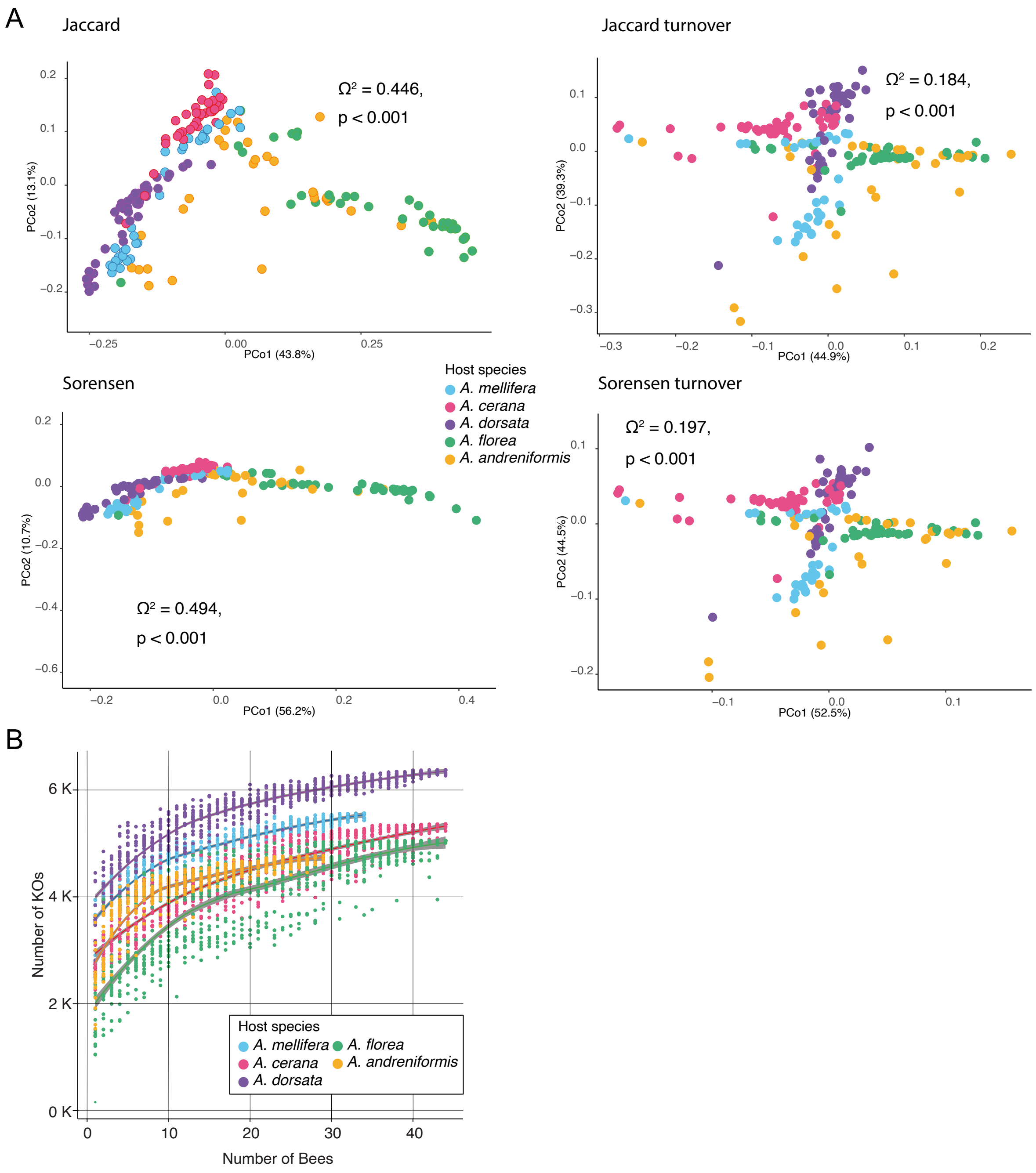


**Supplementary Figure S14. Community-level KO diversity.** (A) PCoA plots using various distance metrics on the KO presence-absence matrix with the result of PERMANOVA for the effect of host species shown on each panel. (B) Cumulative curves of number KOs vs individuals sampled for each host species.

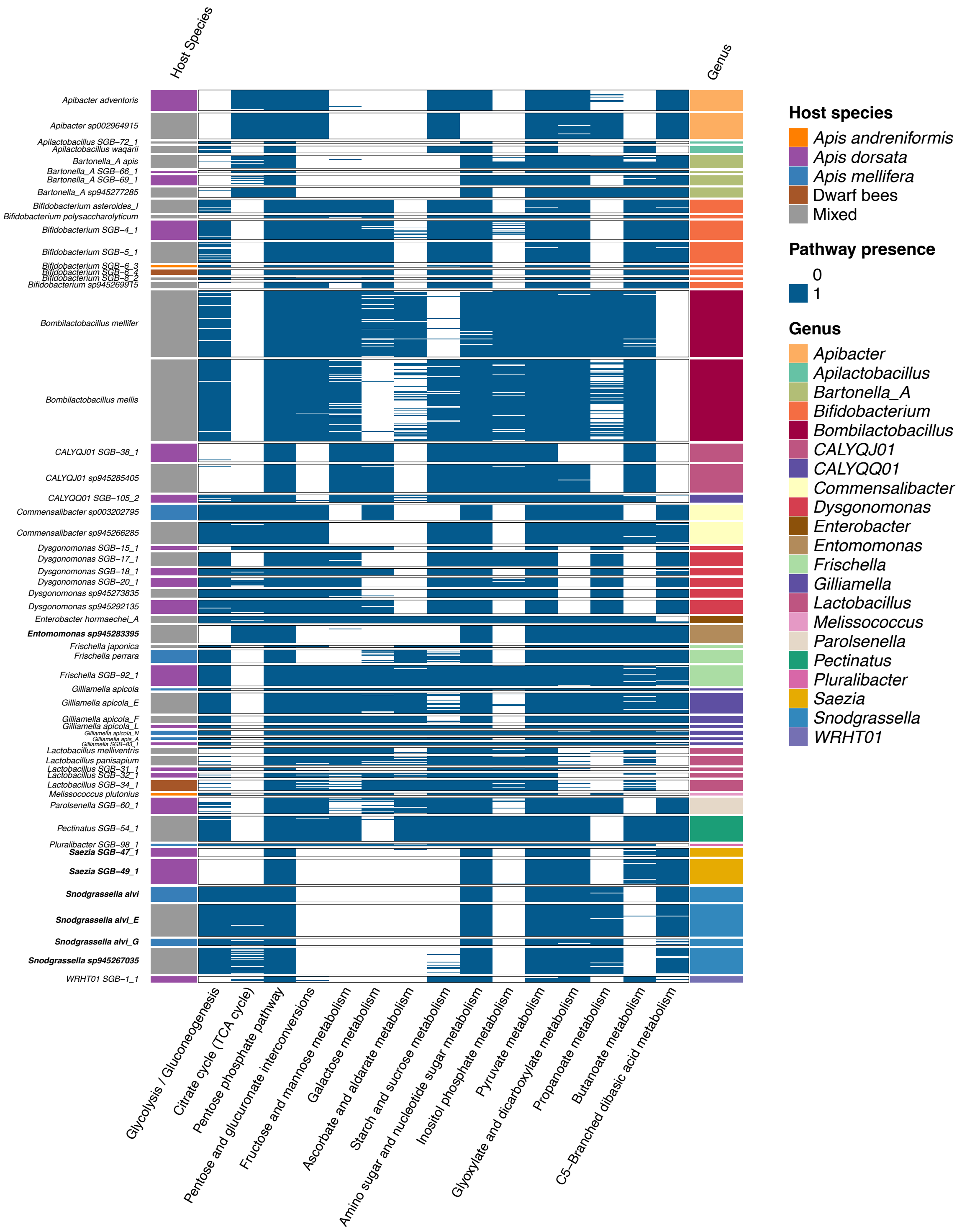


**Supplementary Figure S15. Presence of KEGG Pathways in MAGs.** The heatmap shows high-quality MAGs of different bacterial species as rows and the presence of KEGG pathways shown as columns inferred from the KOs encoded by the respective MAGs using

MinPath. Similar heatmaps including the most prevalent genera and several other pathways are available in the Zenodo (https://zenodo.org/doi/10.5281/zenodo.13732977) repository.

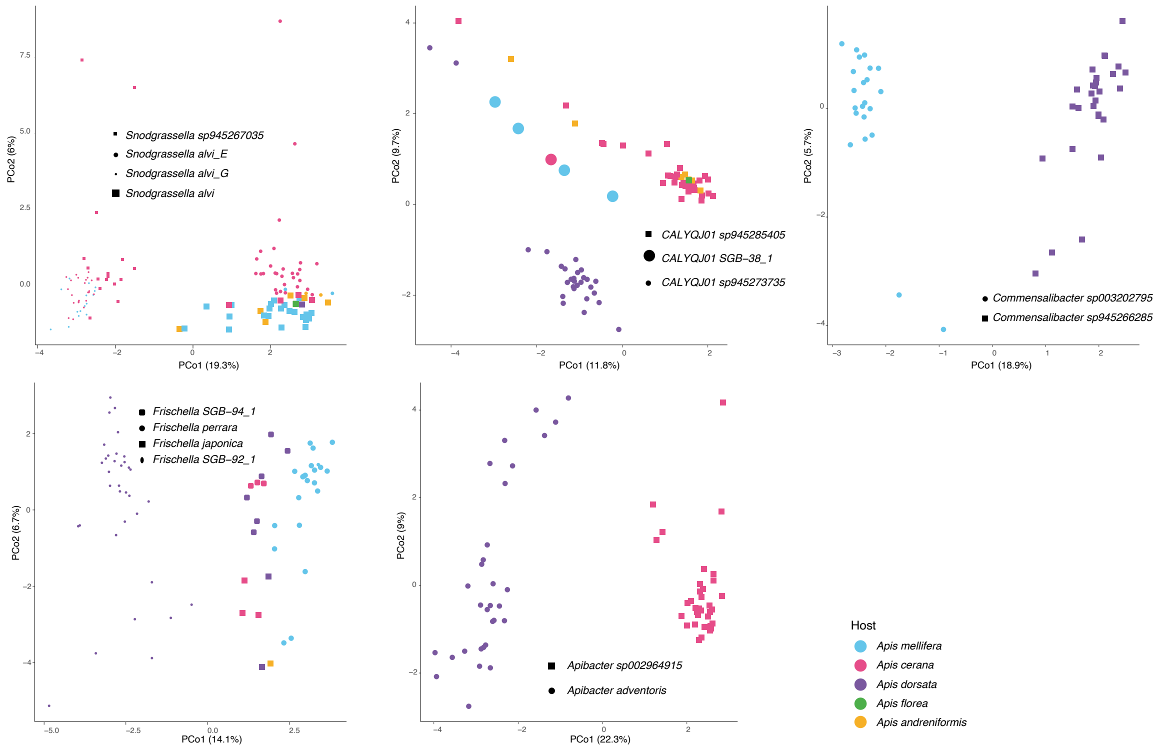


**Supplementary Figure S16. Functional potential of MAGs across bacterial species within the same genus.** PCoA plots of all KOs from MAGs using the Robust-Aitchison distance on the RPKM matrix of bacterial species within a genus with color and shape indicating the host species from which it was recovered and the bacterial species to which it was assigned. Some randomly selected genera and species showing the patterns discussed are displayed.

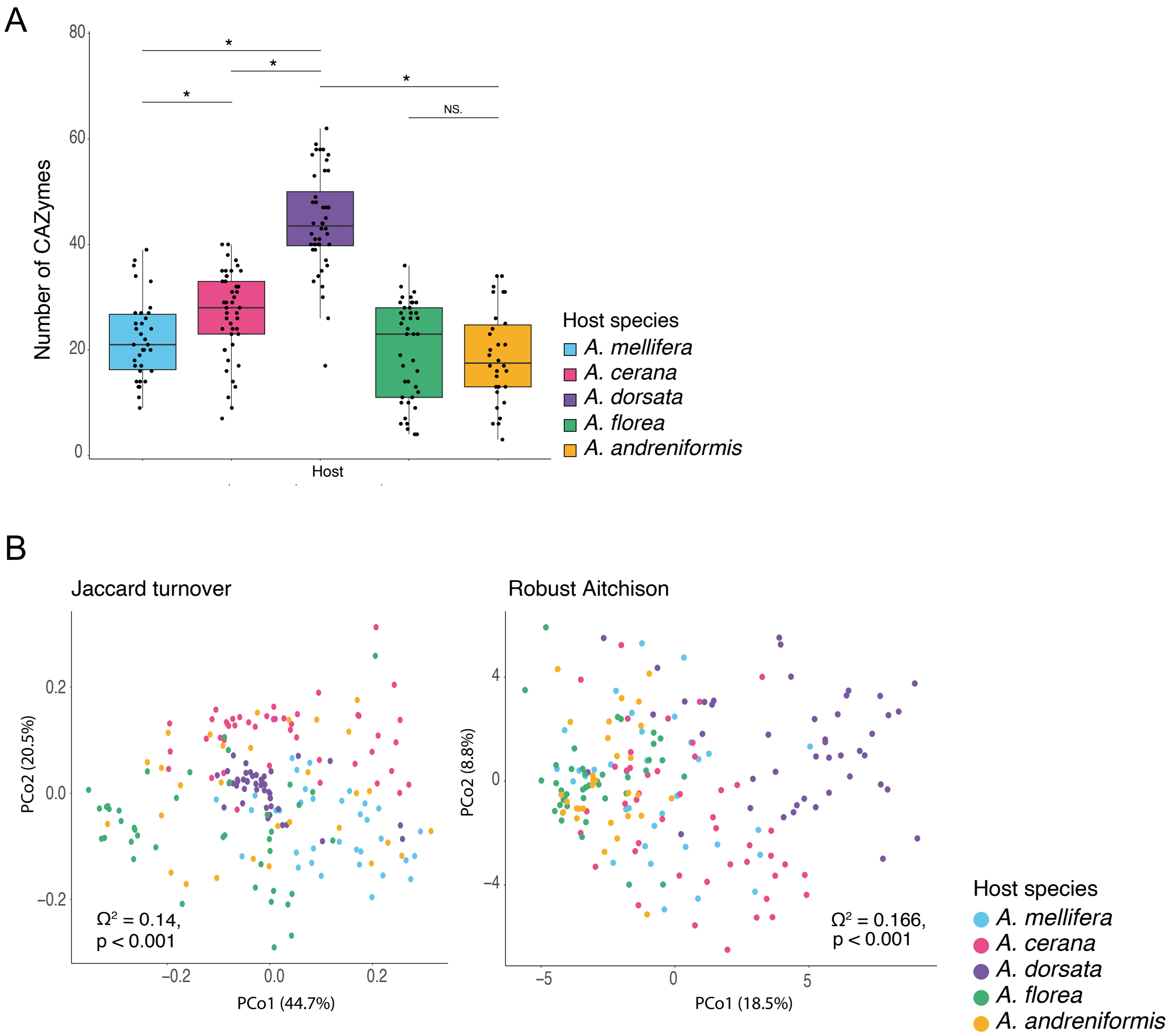


**Supplementary Figure S17. Diversity of CAZymes across honeybee species.** (A) Boxplot showing the number of CAZymes detected per individual bee gut across host species. (B) PCoA plots using various distance metrics on the CAZyme RPKM matrix with the result of PERMANOVA for the effect of host species shown on each panel.

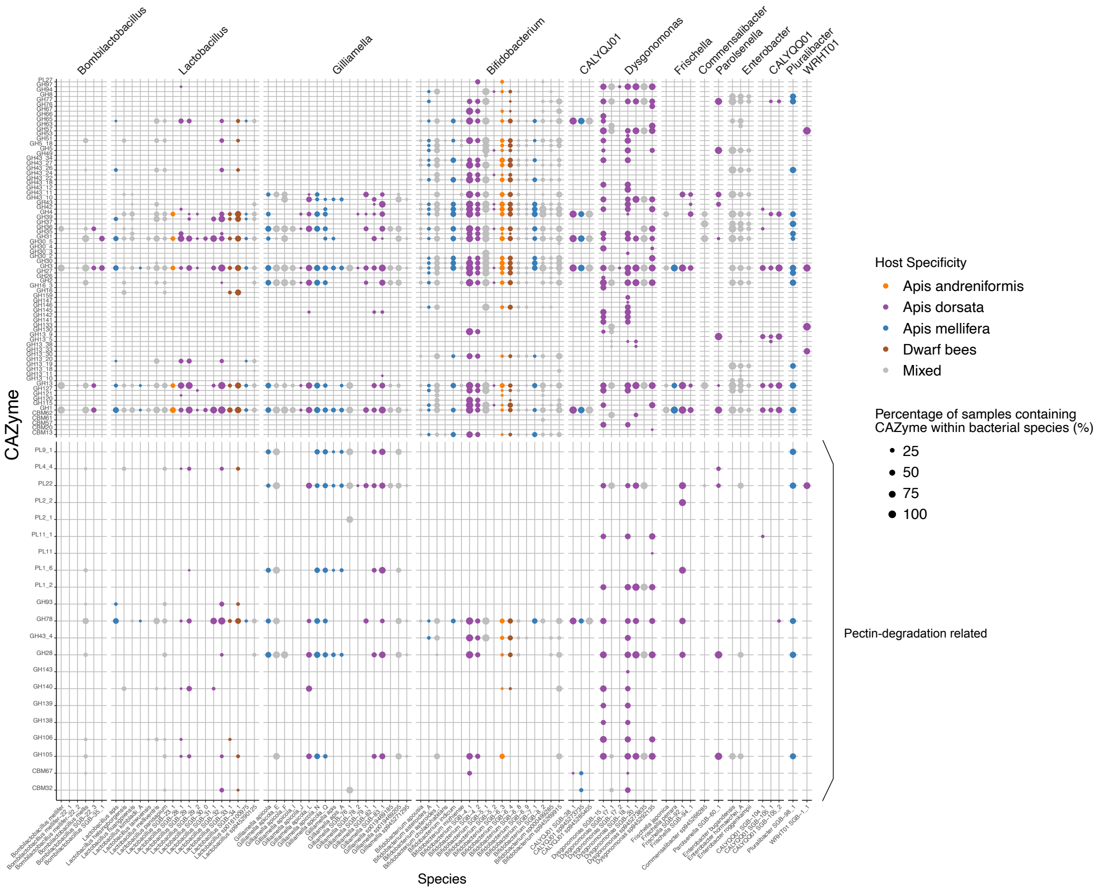


**Supplementary Figure S18. CAZyme families detected in each bacterial species.** Dotplot showing the prevalence (percentage) of CAZymes in each bacterial species. Calculated as the ratio between the number of samples in which the CAZyme is detected on MAGs from the focal bacterial species and the number of samples in which the species was detected. Since this includes several medium-quality MAGs with completeness of only over 50%, CAZymes may not be detected in all MAGs of the species, even if present. The color represents the host species in which the respective bacterial species are found. If it is not a host-specific species, it is marked as mixed. Genera containing at least 5 CAZymes and found in at least 5 samples are included in the plot above (excluding *Escherichia*, *Klebsiella* and *Pantoea*).

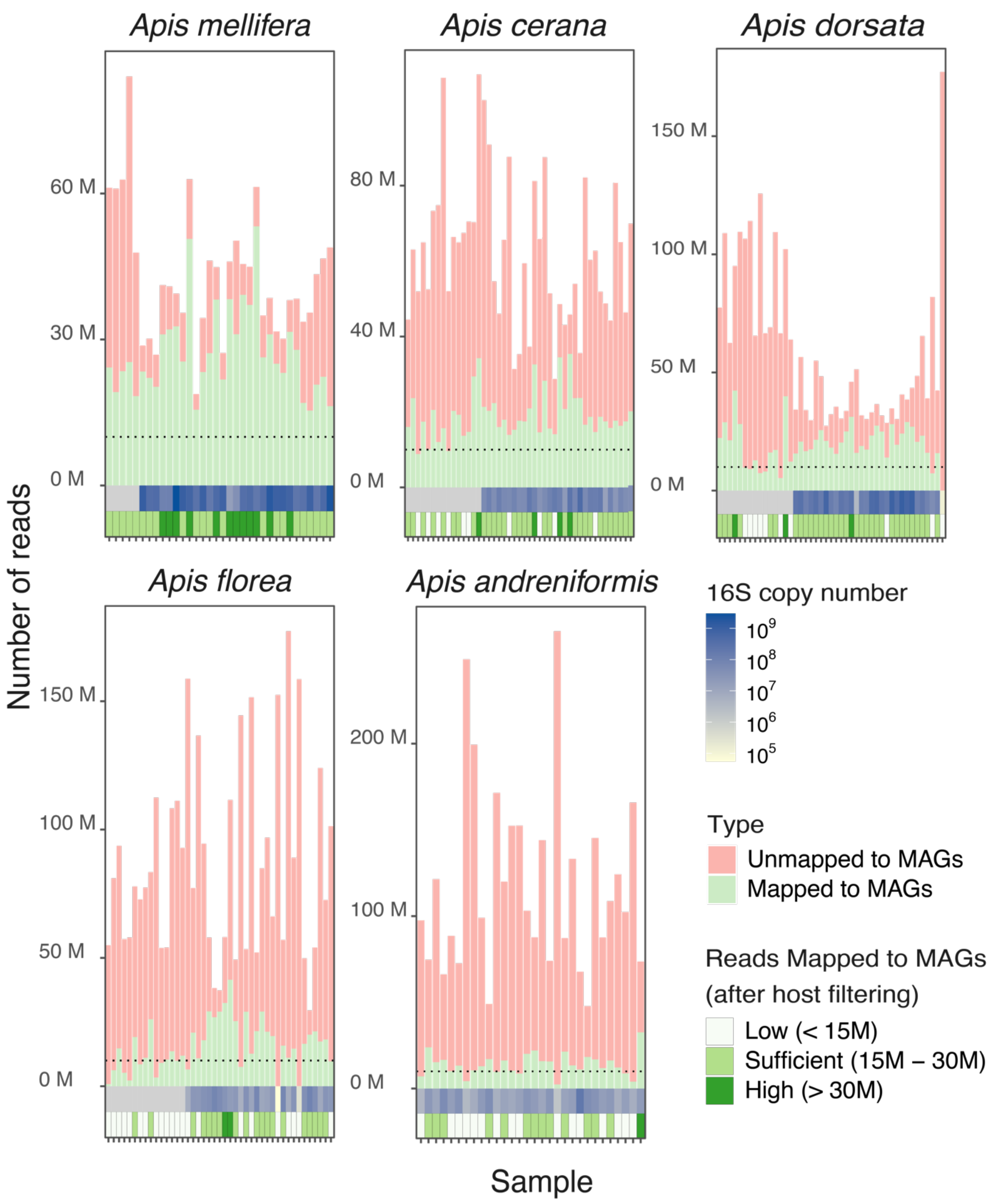


**Supplementary Figure S19. Read mapping to recovered MAGs.** The total number of reads (R1 + R2) from each sample, with the colors showing the number of reads mapped to the MAGs database upon directly mapping trimmed and quality-filtered reads. The dotted line in each panel is drawn at 10M. The tiles below indicate the qPCR copy number using 16S rRNA universal primers (qPCR data not available for samples from India), and the bottom-most tiles show the extent of reads mapped to MAGs after excluding reads mapped to the host genome database.

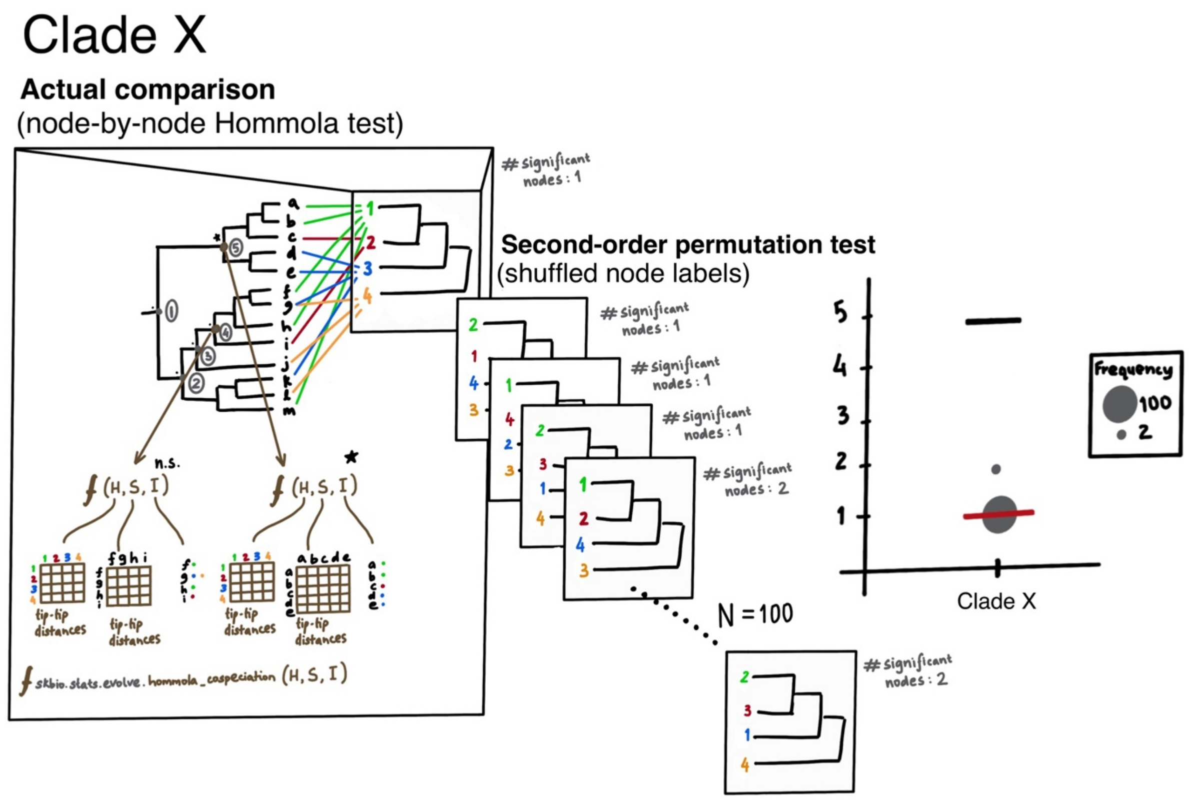


**Supplementary Figure S20. Schematic of co-diversification testing approach.** This schematic describes the approach used to test for co-diversification. For a given clade (in the case of our trees, genus), the nodes that pass filtering thresholds (eg., here, at least 4 MAGs and at least 3 host species represented) are identified. The Hommola cospeciation test (as implemented in the Python package skbio) is applied to each identified node in the tree, considering, in each case, the host matrix (H) and symbiont matrix (S) containing tip-tip distances from phylogeny and the host-symbiont interactions list (I). Subsequently, the number of significant nodes in the tree can be found. Next, to establish a false-discovery (FDR) rate, an expectation for the number of significant nodes can be computed using second-order permutation tests with tip labels of the host tree shuffled in different permutations. Using this, an FDR can be established to indicate whether the significant node(s) identified in the actual comparison is(are) likely to be false positive(s). The black horizontal line on the final plot on the right represents the total number of nodes tested, the grey circles are the number of significant nodes across second-order permutation tests with their size indicating the frequency, and the horizontal red line is the number of significant nodes in the actual comparison.

**Supplementary Data**

**Supplementary Data 1.** (separate file: Supplementary-Data-1.xlsx)

Sample collection information and metadata.

**Supplementary Data 2.** (separate file: Supplementary-Data-2.xlsx)

Metadata about MAGs including fields (in bold) required by the MIMAG standards.

**Supplementary Data 3.** (separate file: Supplementary-Data-3.xlsx)

Sequencing depth and Host and MAG database mapping information.

**Supplementary Data 4.** (separate file: Supplementary-Data-4.xlsx)

Result of multivariate analyses (PERMANOVA test) applied on strain-level profiles for each bacterial species. Column n displays the number of samples in which the species was detected with enough coverage to test for the effect of host species or country on strain-level in a PERMANOVA analysis and the p-value and omega squared values for the respective tests are shown. Host specificity is based on the Rohde's index if strain-level comparison is not available. Values < 1 for which there were not enough samples with good enough coverage for strain-level comparison are considered generalist if there is enough coverage, they are labelled specialist if detected in only one host species, and shared (strain-level specialist) if found in more than one host species but the PERMANOVA test is significant for effect of host species.

**Supplementary Data 5.** (separate file: Supplementary-Data-5.xlsx)

All nodes in original trees identified by hommola test as significant ($p < 0.05$) with the line separating those that are detected with strict thresholds (above the line). sym_subtree displays a Newick string of the subtree at the node being considered. host_tip span is the number of host species represented in the subtree, sym_tips_count the number of MAGs in the subtree.

**Supplementary Data 6.** (separate file: Supplementary-Data-6.xlsx)

Number of nodes tested and found significant under various thresholds. "# significant" is the number of nodes under the respective threshold, "median # significant" is the median number of significant nodes in (N=100) second-order permutation test comparisons with randomly shuffled host tip labels and "SD # significant" is the standard deviation.

**Supplementary Data 7.** (separate file: Supplementary-Data-7.xlsx)

Tip-tip distance between MAGs of the genus in nucleotide tree constructed using 120 core genes (bac120 as identified in GTDB-tk).
